# Kinetics of Hypoglycemia in Diabetes Patients Informs Development of New Modes of Glucagon Therapy

**DOI:** 10.64898/2026.06.05.729994

**Authors:** Jason Li, Ceara Byrne, Jia Y. Liang, SeJun Lee, Mille Kronborg Lyhne, Anika Meng, Susan Rui Ling, Shiyi Li, Aaron Lopes, Parmiss Khosravi, Colin Cotter, Yuyan Su, Ethan D’Orio, Johannes Josef Fels, Jacob W. Coffey, Alison Hayward, Andreas Vegge, Ulrik Rahbek, Stephen T. Buckley, Robert Langer, Giovanni Traverso

**Affiliations:** Division of Gastroenterology, Hepatology and Endoscopy, Brigham and Women’s Hospital, Harvard Medical School; Boston, MA 02115, USA; David H. Koch Institute for Integrative Cancer Research, Massachusetts Institute of Technology; Cambridge, MA 02139, USA; Broad Institute of MIT and Harvard; Cambridge, MA 02139, USA; Department of Mechanical Engineering, Massachusetts Institute of Technology; Cambridge, MA 02139, USA; Research & Early Development, Novo Nordisk A/S, Copenhagen, Denmark; Faculty of Health and Medical Sciences, University of Copenhagen; Department of Microbiology and Immunology, Peter Doherty Institute for Infection and Immunology, University of Melbourne, Melbourne, VIC, 3000 Australia; Division of Comparative Medicine, Massachusetts Institute of Technology; Cambridge, MA 02139, USA; Department of Chemical Engineering, Massachusetts Institute of Technology, Cambridge; MA 02139, USA

## Abstract

Insulin therapy revolutionized the care of patients with diabetes starting ∼100 years ago, yet insulin-induced hypoglycemia remains a serious life-threatening complication of insulin therapy. Glucagon is a highly effective treatment; however current dosage forms remain under-utilized due to poor patient compliance. The development of improved and situation-specific glucagon therapies remains challenging due to the poor drug stability and incomplete knowledge of the kinetics of different hypoglycemic events. Thus, we analyzed continuous glucose monitor (CGM) data from 1135 patients with type 1 diabetes (T1D) representing 246.18 patient years. We show that a surprisingly large proportion of hypoglycemic episodes (20-30%) are follow-on events resulting from under-treatment of prior events, and that the average duration of independent hypoglycemic events can last up to 79 to 108 minutes. We further show that the kinetics of hypoglycemic onset and persistence varies significantly by patient history, severity, time of occurrence. Guided by these findings, we recognize the opportunity to develop high-density, readily-soluble, and thermostable (ReST) solid glucagon formulations, and painless application-specific microneedle-patches that are in line with the timing needs of T1D patients who are awake and asleep. Thus, we demonstrate (1) on-demand patches for rapid prevention or treatment of mild hypoglycemia during the day, and (2) enzyme-driven hypoglycemia-responsive patches supporting autonomous glucagon release during the night. We show excellent in vitro glucagon stability, loading, and release kinetics of both systems and demonstrate their ability to treat hypoglycemia in diabetic animals. The engineering of these delivery systems demonstrates the potential of human CGM data and solid glucagon formulations to enable new modes of glucagon therapy, thereby expanding the clinical role of glucagon beyond the emergency setting.

## Introduction

Hypoglycemia is a life-threatening condition that is prevalent in people with diabetes due to over-administration of insulin^1–3^. This acute condition can cause debilitating mental or physical symptoms and may progress to loss of consciousness or death^4^. Thus, hypoglycemia remains the most serious acute complication among patients with diabetes and often limits the use of intensive insulin therapy resulting in an increased risk of long-term diabetic complications^1,2^.

While the importance of timely treatment of severe and symptomatic hypoglycemia has long been recognized, it is becoming evident that rapid treatment of mild asymptomatic events is vital to prevent dysregulation of the body’s counterregulatory hormone response and progressive blunting of the brain’s response to low glucose levels; both of which can lead to a vicious cycle of recurrent hypoglycemia^2,5–7^. Preventing mild hypoglycemia progression to severe hypoglycemia is also critical as recent studies have shown that patients that experience severe hypoglycemia are at greater risk of all-cause mortality in the short term after the hypoglycemic episode^8^. However, approved glucagon treatments for mild hypoglycemia (e.g., mini-dose glucagon therapy) are not available despite clinical evidence suggesting their effectiveness and user preference^9–12^. Furthermore, the use of glucagon for the treatment of severe hypoglycemia remains under-utilized due to the pain and complexity associated with current glucagon interventions^13,14^. Prophylactic treatments for hypoglycemia and counter-regulatory closed-loop systems offer significant benefits for improving hypoglycemia treatment rates, however FDA-approved therapies of this nature do not exist. Thus, there is considerable interest in expanding the use of glucagon beyond emergency rescue treatment of severe hypoglycemia^15^.

To date, the development of targeted and effective glucagon treatments for hypoglycemic conditions beyond rescue treatment of severe hypoglycemia is hindered by our incomplete knowledge of the temporal trajectory of these events, particularly in the home setting, and the availability of painless and ready-to-administer thermostable glucagon formulations^14^. While many aspects of hypoglycemic events are well characterized in various patient populations (e.g., prevalence, cumulative time in hypoglycemia, and variations in diurnal occurrence), little is known about the kinetics of hypoglycemia onset, duration, and time-to-treatment for individual events^7,16–19^. Such granular data is vital for engineering novel glucagon treatments as it enables the development of patient-centric and situation-specific treatment offerings, and clinically meaningful engineering design requirements.

The insoluble nature of glucagon and its tendency to rapidly degrade and form insoluble fibrils in solutions has prevented the development of new glucagon formulations for many years^20^. The recent introduction of solution-stable glucagon formulations improves upon earlier emergency glucagon rescue kits since they are thermostable during storage and ready for administration through injection^21,22^. However, pain and fear associated with needle injections remain a significant clinical challenge resulting in suboptimal use of an otherwise effective medication, particularly in non-emergent situations^2,23^. Intranasal glucagon formulations circumvent this limitation; however, drawbacks include painful respiratory side-effects, delayed resolution to hypoglycemia, and inaccurate dose administration which precludes use outside of emergencies and for treatment of mild hypoglycemia^24–26^. Furthermore, the introduction of these ready-to-use injectable or inhalable glucagon formulations has not been successful at expanding the use of glucagon in patients^27^, and their low drug density limits efficient loading and delivery via more patient-friendly ingestible or transdermal delivery devices^28–30^. Thus, the insoluble and unstable nature of glucagon continues to stifle the development and availability of painless and well-tolerated glucagon treatments for a variety of hypoglycemic conditions beyond severe hypoglycemia.

In this work, we performed a retrospective analysis of continuous glucose monitoring (CGM) data from human subjects with type 1 diabetes (T1D) to identify the trajectories of hypoglycemic events based on severity and time-of-occurrence; and derive clinically meaningful design parameters (e.g., *pre-hypoglycemic duration*, *mild hypoglycemic duration preceding severe hypoglycemia*, and *time to treatment*) to guide the development of new glucagon interventions. We further identify high-density readily soluble and thermostable (ReST) solid glucagon formulations that enable the storage and administration of glucagon in the dry form. We leverage these findings to develop two transdermal glucagon delivery systems: A space-efficient, painless, and convenient microneedle patch for accurate mini-dose glucagon treatment of mild hypoglycemia and scalable for larger doses, and an enzymatic glucose-responsive wearable patch that provides automated and glucose-specific prophylactic treatment of nocturnal hypoglycemia throughout the night. These implementations demonstrate the potential of the high-density ReST glucagon formulations to enable new modes of glucagon therapy, thereby expanding the clinical role of glucagon beyond the emergency setting, facilitating more widespread management of hypoglycemia, and improving the glycemic management of people with diabetes.

## Results

### Trajectories of hypoglycemic events T1D

The aggregated dataset is comprised of 246.18 patient-years of CGM data from 5 publicly available clinical trial datasets and included 1135 subjects (Table 1)^16,31–35^. Patient enrollment criteria suggests that 909 subjects from 4 studies represent the *general type 1 diabetic population* while the remaining 226 subjects are considered representative of T1D patients with *well-controlled diabetes* with no recent history of hypoglycemia. A total of 158344 distinct hypoglycemic events were identified in the dataset, each represented by a CGM trace comprised of a *pre-hypoglycemic period* and a *hypoglycemic period* during which blood glucose levels remain below 70 mg/dL (Fig. 1a).

**Fig. 1.**
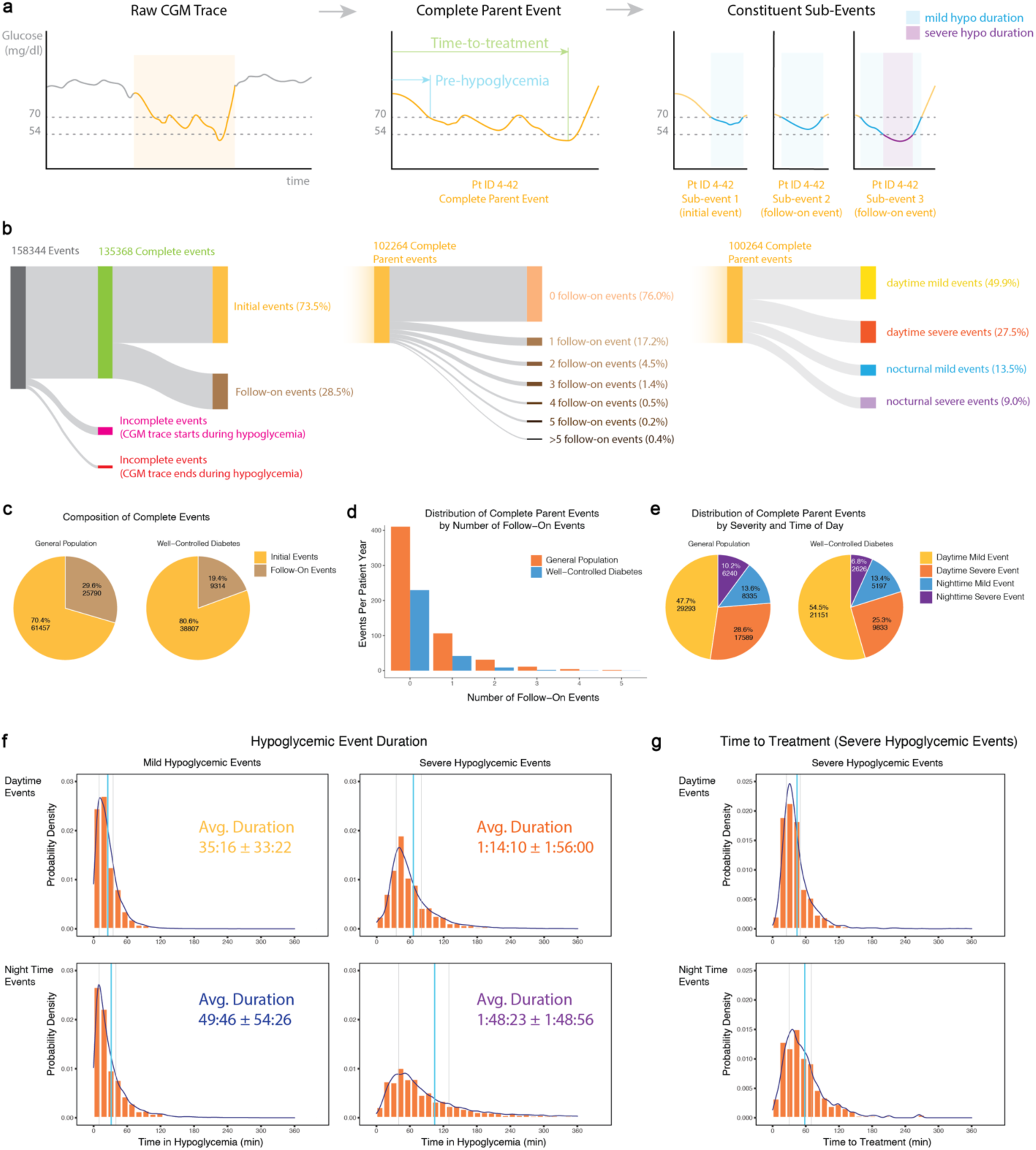
Characteristics of hypoglycemic events. a) Extraction and parameterization of hypoglycemic events from raw CGM trace. b) Sankey diagrams showing the classification of hypoglycemic events. c) Proportion of complete initial and follow-on events in the general and well-controlled T1D populations. d) Incidence of parent events by several associated follow-on events. e) Proportion of complete parent events by hypoglycemic severity and time of occurrence. f) Probability density of hypoglycemia durations by hypoglycemia severity and time of day. Average duration (blue vertical line) and 25th and 75th percentiles (grey vertical line) are indicated in each plot. g) Probability density of time-to-treatment following the onset of hypoglycemia. Average time (blue vertical line) and 25th and 75th percentiles (grey vertical line) are indicated in each plot.

**Table 1.** Participant Characteristics.

|  | All Subjects | Well-controlled T1D | General T1D | Alep. | Buck. | Chase | Tamb. | Wein. |
| --- | --- | --- | --- | --- | --- | --- | --- | --- |
| Number of Participants - no. | 1115 | 226 | 889 | 226 | 30 | 207 | 451 | 201 |
| Diabetes Type | Type 1 | Type 1 | Type 1 | Type 1 | Type 1 | Type 1 | Type 1 | Type 1 |
| Sex - no. (%) |  |  |  |  |  |  |  |  |
| Female | 563 (50.9) | 112 (49.6) | 451 (50.7) | 112 (49.6) | 12 (40) | 96 (46.4) | 248 (55) | 95 (47.3) |
| Male | 544 (49.1) | 114 (50.4) | 438 (49.3) | 114 (50.4) | 18 (60) | 111 (53.6) | 203 (45) | 106 (52.7) |
| Age - yr |  |  |  |  |  |  |  |  |
| average (yr) | - | - | - | 44 +/-14 | 11.2+/-4.1 | 12.5+/-2.8 | 25 +/- 15.8 | - |
| range (yr) | 3-60+ | - | - | 19-78 | 3-17 | 7-18 | 8+ | 60+ |
| Race / ethnicity |  |  |  |  |  |  |  |  |
| White non-hispanic | 981 | 207 | 774 | 207 | - | 172 | 417 | 185 |
| Hispanic or Latino | 35 | 9 | 26 | 9 | - | 5 | 15 | 6 |
| Black / African American | 33 | 5 | 28 | 5 | - | 14 | 7 | 7 |
| Asian | 11 | 4 | 7 | 4 | - | 4 | 2 | 1 |
| American Indian / Alaskan Native | 2 | 0 | 2 | 0 | - | 0 | 1 | 1 |
| Native Hawaiian / Pacific Islander | 1 | 0 | 1 | 0 | - | 0 | 1 | 0 |
| Multiple Ethnicities | 15 | 0 | 15 | 0 | - | 10 | 5 | 0 |
| Other / Unknown | 7 | 1 | 6 | 1 | - | 2 | 3 | 1 |
| Glycated hemoglobin HbA1c - % | - | 7.1 ± 0.6 | - | 7.1 ± 0.6 | 7.1 ± 0.6 | 8.0 ± 0.9 | 322 (7-10% A1c)<br>129 (<7% A1c) | - |
| Body-mass index (kg/m2) | 24.3 ± 5.0 | 27.3 ± 4.4 | 23.5 ± 4.8 | 27.3 ± 4.4 | 19.8 ± 3.5 | 20.6 ± 3.7 | 23.5 ± 4.4 | 26.9 ± 4.7 |
| Body-mass index z score - no. (%) |  |  |  |  |  |  |  |  |
| < -0.5 | 33.36% | 34.51% | 33.07% | 34.51% | 30.00% | 38.16% | 31.26% | 32.34% |
| -0.5 to 0.5 | 30.94% | 38.94% | 28.91% | 38.94% | 20.00% | 31.88% | 40.13% | 1.99% |
| >0.5 | 35.07% | 26.55% | 37.23% | 26.55% | 26.67% | 29.95% | 28.60% | 65.67% |
| no data | 0.63% | 0.00% | 0.79% | 0.00% | 23.33% | 0.00% | 0.00% | 0.00% |
| Hypoglycemia history |  |  |  |  |  |  |  |  |
| one or more episodes of severe hypoglycemia during previous 6 mo - no. (%) | - | No history of severe hypoglycemia | No Screening | No history of severe hypoglycemia | No Screening | No Screening | No Screening | No Screening |
| insulin administration - no. (%) |  |  |  |  |  |  |  |  |
| pump | - | 98.67% | - | 98.67% | - | 53.14% | 81.37% | 58.71% |
| multiple daily injections | - | 1.33% | - | 1.33% | - | 46.86% | 18.63% | 41.29% |
| **Study design** |  |  |  |  |  |  |  |  |
| Participant enrollment period | 2-26 weeks | 26 weeks (6 months) | 2-26 weeks | 26 weeks (6 months) | 13 weeks | 26 weeks (6 months) | 26 weeks (6 months) | 2 weeks |
| CGM Sample Frequency (min) | 5 | 5 | 5 | 5 | 5 | 5 | 5 | 5 |
| CGM Device | Multiple CGM Devices | Dexcom G4 | Multiple CGM Devices | Dexcom G4 | FreeStyle Navigator | Medtronic CGMS, Cygnus GlucoWatch G2 Biographer | FreeStyle Navigator, Dexcom SEVEN, Medtronic Paradigm | Dexcom SEVEN PLUS |

### Incidence Rate

Complete hypoglycemic events were common among the *general* and *well-controlled* T1D patient populations with incidence rates of 803 and 350 events per patient-year respectively (Table 2). Independent hypoglycemic events are perhaps a more clinically relevant metric of incidence given that *follow-on events* are likely a direct result of inadequate treatment of a prior event. Therefore, we took a more holistic approach in our analysis by associating any follow-on events with the independent hypoglycemic events that closely precede them. Thus, the incidence rate of *parent events* in the general population was 566 events per patient-year (70.4% of all *complete events*), and 282 events per patient-year in the well-controlled diabetes population (80.6% of all *complete events*) (Fig 1c). Patients with well-controlled T1D experience 43.6% to 49.9% fewer *parent events* compared to the general T1D population and ∼10% fewer follow-on events. A surprisingly high proportion (i.e., 24.0%) of parent events are associated with one or more follow-on events indicating that under-treatment of hypoglycemia is fairly common (Fig. 1b, d).

**Table 2.**
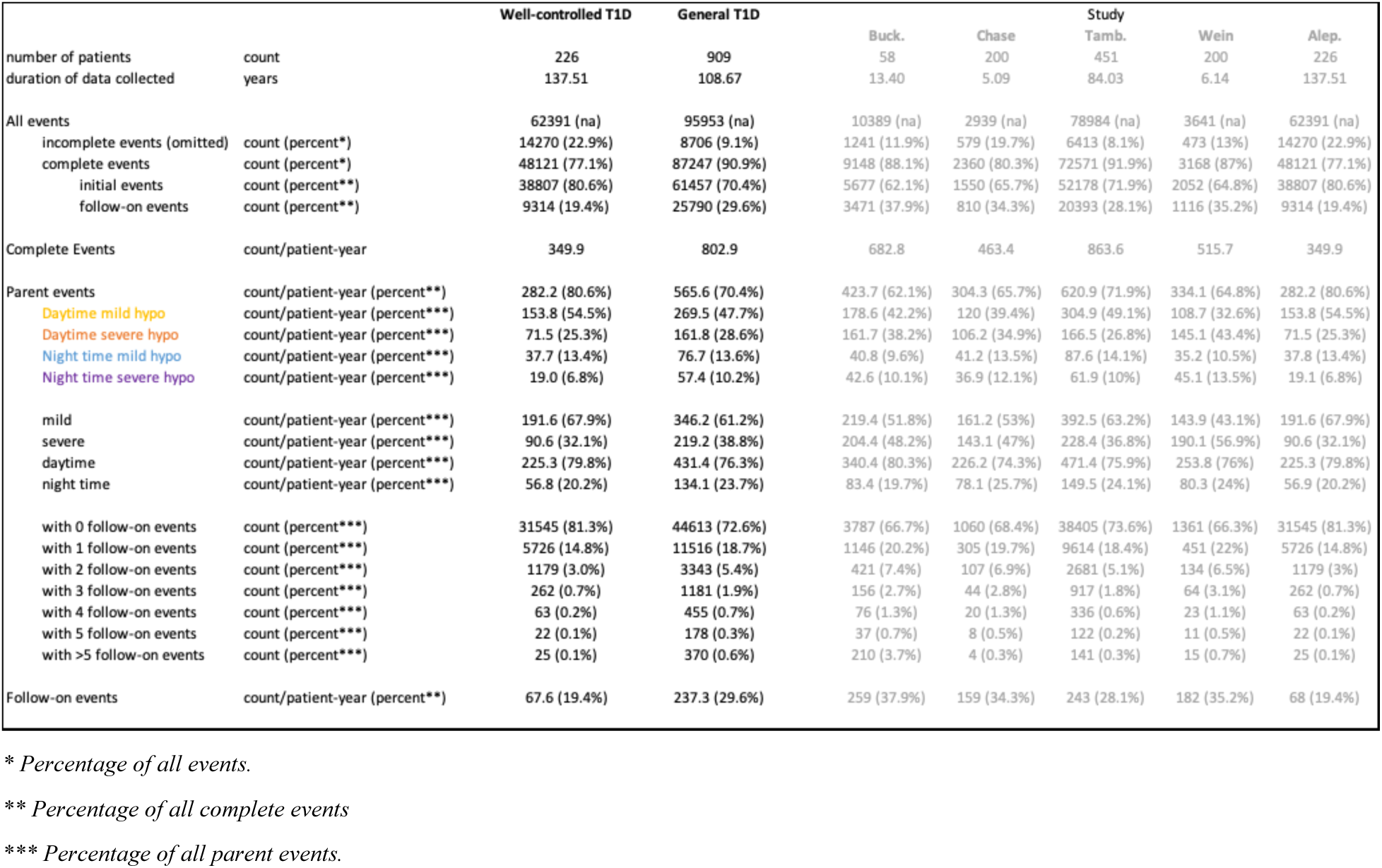
Incidence of Hypoglycemic Events.

Segmentation of parent hypoglycemic events by severity show that 61-68% of these instances are mild hypoglycemia (between 54 and 70 mg/dL), with the remaining 32-39% of events being severe cases of hypoglycemia (<54 mg/dL) (Fig. 1b, e). The incidence rate of *severe parent events* in the general T1D population is on average 219 events per patient-year which is 1.5 to 2.5-fold higher than the 91 *severe parent events* per patient-year observed in people with well-controlled T1D. These results show that the incidence of hypoglycemia is very common both during the day and at night, especially for mild cases. These rates are lower than previously reported (i.e., 0.25-0.60 versus ∼1 events per patient per day^31,36^), likely due to treatment of an initial and subsequent follow-on events as discrete events and differences in the hypoglycemic thresholds used in the analysis.

Classification of parent hypoglycemic events by time of occurrence reveal that 76-80% of events occur during the day, with the balance 20-24% of events occurring at night (Fig. 1e). A spike in both mild and severe hypoglycemic parent events was observed at 12:00 (noon) and 18:00 hours, presumably due to meals, as well as at midnight (00:00) (Fig. S2). This temporal distribution is in agreement with previous studies^18,37^.

### Hypoglycemia onset: Rate and time in pre-hypoglycemia

The pre-hypoglycemic period represents a window of opportunity to prevent impending hypoglycemia following the administration of excessive insulin and was found to differ based on the severity and time of occurrence (Table 3, Fig. S3). On average, this duration was 14.3% to 17.3% shorter for severe events compared to mild events and was accompanied by faster rates of blood glucose decline (−2.54 to -5.2 mg/dL/min) compared to mild events (−0.88 to -1.05 mg/dL/min).

**Table 3.** Characteristics of Parent Hypoglycemic Events.

|  |  | Well-controlled T1D | General T1D | Statistical Significance (mean values) |  |  |  |  |
| --- | --- | --- | --- | --- | --- | --- | --- | --- |
|  |  |  |  | general vs.<br>well-controlled | vs. day mild | vs. day severe | vs. night mild | vs. night severe |
| Rate of hypoglycemia onset |  |  |  |  |  |  |  |  |
| Daytime mild hypo | mean ± SD [mg/dl/min] | -1.02 ± 2.07 | -1.05 ± 1.98 | P = 0.01058 | na | P < 2.2e-16 | P < 2.2e-16 | P < 2.2e-16 |
| Daytime severe hypo | mean ± SD [mg/dl/min] | -1.44 ± 1.37 | -5.2 ± 176.2 | P < 2.2e-16 | P < 2.2e-16 | na | P < 2.2e-16 | P < 2.2e-16 |
| Night time mild hypo | mean ± SD [mg/dl/min] | -0.84 ± 0.85 | -0.88 ± 0.87 | P = 0.1278 | P < 2.2e-16 | P < 2.2e-16 | na | P < 2.2e-16 |
| Night time severe hypo | mean ± SD [mg/dl/min] | -1.19 ± 1.05 | -2.54 ± 47.96 | P = 4.767e-10 | P < 2.2e-16 | P < 2.2e-16 | P < 2.2e-16 | na |
| Pre-hypoglycemia Duration |  |  |  |  |  |  |  |  |
| Daytime mild hypo | mean ± SD (median, IQR) [h:m:s] | 46:57 ± 38:32 (35:00) | 1:14:13 ± 56:06 (1:00:00) | P < 2.2e-16 | na | P < 2.2e-16 | P = 1.369e-5 | P = 1.577e-11 |
| Daytime severe hypo | mean ± SD (median, IQR) [h:m:s] | 47:32 ± 36:55 (40:00) | 1:01:25 ± 46:52 (50:00) | P < 2.2e-16 | P < 2.2e-16 | na | P = 6.408e-4 | P = 0.7667 |
| Night time mild hypo | mean ± SD (median, IQR) [h:m:s] | 44:58 ± 41:28 (30:00) | 1:19:04 ± 1:08:39 (55:00) | P < 2.2e-16 | P = 1.369e-5 | P = 6.408e-4 | na | P = 0.03092 |
| Night time severe hypo | mean ± SD (median, IQR) [h:m:s] | 47:08 ± 40:38 (35:00) | 1:07:06 ± 59:12 (50:00) | P < 2.2e-16 | P = 1.577e-11 | P = 0.7667 | P = 0.03092 | na |
| Duration of mild hypo preceding severe hypo |  |  |  |  |  |  |  |  |
| Daytime severe hypo | mean ± SD (median, IQR) [h:m:s] | 16:48 ± 13:56 (15:00) | 15:59 ± 18:15 (10:00) | P < 2.2e-16 |  |  |  | P < 2.2e-16 |
| Night time severe hypo | mean ± SD (median, IQR) [h:m:s] | 25:00 ± 23:01 (20:00) | 25:14 ± 31:53 (15:00) | P = 1.735e-06 |  | P < 2.2e-16 |  |  |
| Hypoglycemia Duration |  |  |  |  |  |  |  |  |
| Daytime mild hypo | mean ± SD (median, IQR) [h:m:s] | 26:03 ± 20:13 (20:00) | 35:16 ± 33:22 (25:00) | P < 2.2e-16 | na | P < 2.2e-16 | P = 3.175e-16 | P < 2.2e-16 |
| Daytime severe hypo | mean ± SD (median, IQR) [h:m:s] | 38:51 ± 26:10 (30:00) | 1:14:10 ± 1:56:00 (47:00) | P < 2.2e-16 | P < 2.2e-16 | na | P < 2.2e-16 | P < 2.2e-16 |
| Night time mild hypo | mean ± SD (median, IQR) [h:m:s] | 32:06 ± 31:14 (25:00) | 49:56 ± 54:26 (30:00) | P < 2.2e-16 | P = 3.175e-16 | P < 2.2e-16 | na | P < 2.2e-16 |
| Night time severe hypo | mean ± SD (median, IQR) [h:m:s] | 48:39 ± 39:23 (35:00) | 1:48:23 ± 1:48:56 (1:10:00) | P = 2.871e-12 | P < 2.2e-16 | P < 2.2e-16 | P < 2.2e-16 | na |
| Duration to first glucagon treatment |  |  |  |  |  |  |  |  |
| Daytime severe hypo | mean ± SD (median, IQR) [h:m:s] | 37:29 ± 19:09 (35:00) | 49:34 ± 42:36 (40:00) | P = 1.46e-12 |  | na |  | P = 7.204e-15 |
| Night time severe hypo | mean ± SD (median, IQR) [h:m:s] | 38:47 ± 24:01 (45:00) | 57:35 ± 50:43 (45:00) | P = 0.8339 |  | P = 7.204e-15 |  | na |
| Duration to last glucagon treatment |  |  |  |  |  |  |  |  |
| Daytime severe hypo | mean ± SD (median, IQR) [h:m:s] | 53:29 ± 33:40 (35:00) | 54:14 ± 52:43 (40:00) | P = 0.0122 |  | na |  | P = 0.8077 |
| Night time severe hypo | mean ± SD (median, IQR) [h:m:s] | 54:12 ± 35:09 (45:00) | 1:03:00 ± 1:05:59 (45:00) | P = 0.5817 |  | P = 0.8077 |  | na |

Data reported as mean ± SD [hr:min:sec] (median, IQR)

The pre-hypoglycemic duration for mild nocturnal hypoglycemic events was 6.5% longer than mild daytime events (p = 1.369e^-5^). This may be due to the use of short-acting insulin during the day compared to the use of long-acting insulin before sleep. Notably, no significant difference in the pre-hypoglycemic duration was observed between daytime and nighttime severe events (p = 0.7667). Finally, the average duration of pre-hypoglycemia was found to be 29.2% to 75.8% longer in the general T1D population compared to patients with well-controlled disease (p < 2.2e^-16^).

### Hypoglycemic event duration

The various timings of hypoglycemic events represent multiple windows of opportunity for intervention including the opportunity to mitigate progression of mild to severe hypoglycemia, and the opportunity to provide automated glucagon treatment of severe hypoglycemia. The aggregate CGM traces during hypoglycemia are shown in Figure S3. The average duration of parent hypoglycemic events is dependent on severity and time of occurrence (Table 3). During the day, the average mild hypoglycemic event lasted 35.27 minutes compared to the average severe hypoglycemic event which lasted 74.17 minutes, representing a 110.3% longer duration for severe events (p<2.2e^-16^) (Figure 1f, S4). During the night, the average mild hypoglycemic event lasted 49.77 minutes and the average severe hypoglycemic event lasted 108.38 minutes, representing a 117.8% longer duration for severe events (p<2.2e^-16^). On average, nocturnal hypoglycemic events were longer than daytime events (i.e., 41.1% for mild events, p = 3.175e^-16^; and 46.1% for severe events, p < 2.2e^-16^). Finally, the average duration of each hypoglycemic event (daytime mild, daytime severe, nighttime mild, nighttime severe) was between 35.3% and 108.4% longer in the general T1D population compared to patients with well-controlled disease (p < 2.2e^-16^ for daytime mild, daytime severe, nighttime mild; p=2.871e^-12^ for nighttime severe).

The average time to glucagon treatment of severe hypoglycemia in the general T1D population was 49.57 minutes during the day and 57.58 minutes during the night (p = 7.2e^-15^) (Figure 1g, S5). A trend towards shorter treatment times was observed in patients with well-controlled T1D during the day (37.48 minutes, p = 1.46e^-12^), but this difference was not significant at night (38.78 minutes, p = 0.8339).

The duration of a hypoglycemic event consists of the time it takes for patients to recognize the onset of hypoglycemia, administer an intervention, and the time needed for that intervention to restore euglycemia. While any treatment should ideally rescue a patient from hypoglycemia in as short a timeframe as possible, the reported durations reflect current medical practice and therefore a minimum threshold for which a novel glucagon treatment should strive to restore euglycemia to be considered a viable treatment option.

### In vitro screening and characterization of high-density readily soluble and thermostable (ReST) solid glucagon formulations

A high-throughput formulation screen was performed to identify solid pharmaceutical excipients capable of promoting dissolution of lyophilized glucagon in neutral pH solutions. Among the 141 excipients tested, 10 compounds successfully enhanced the dissolution of glucagon following a 30-minute incubation in pH 7.4 PBS at 37°C (Fig. 2a). These excipients enhanced solution glucagon concentrations by up to 16-fold compared to unformulated glucagon. Many of the top hits (e.g., Benzoic acid and sodium carbonate) promote glucagon dissolution by altering solution pH (Fig. S6). The zwitterionic compound myristyl sulfobetaine (MSB) is notable for its ability to enhance glucagon dissolution in PBS without altering solution pH (Fig. S6). Interestingly, most zwitterionic compounds tested, including those with similar chemical structures to MSB (e.g., lauryl sulfobetaine, octyl sulfobetaine), were unable to promote glucagon dissolution. Further evaluation of the *in vitro* dissolution kinetics of a compacted solid glucagon-MSB formulation demonstrated the ability of MSB to promote rapid and complete dissolution of lyophilized glucagon in a dose dependent manner in neutral pH buffer compared to unformulated drug, which remained largely insoluble (Fig. 2b, 2c). Therefore, readily soluble high-density binary solid formulations containing as much as 80% wt. glucagon is enabled.

**Fig. 2.**
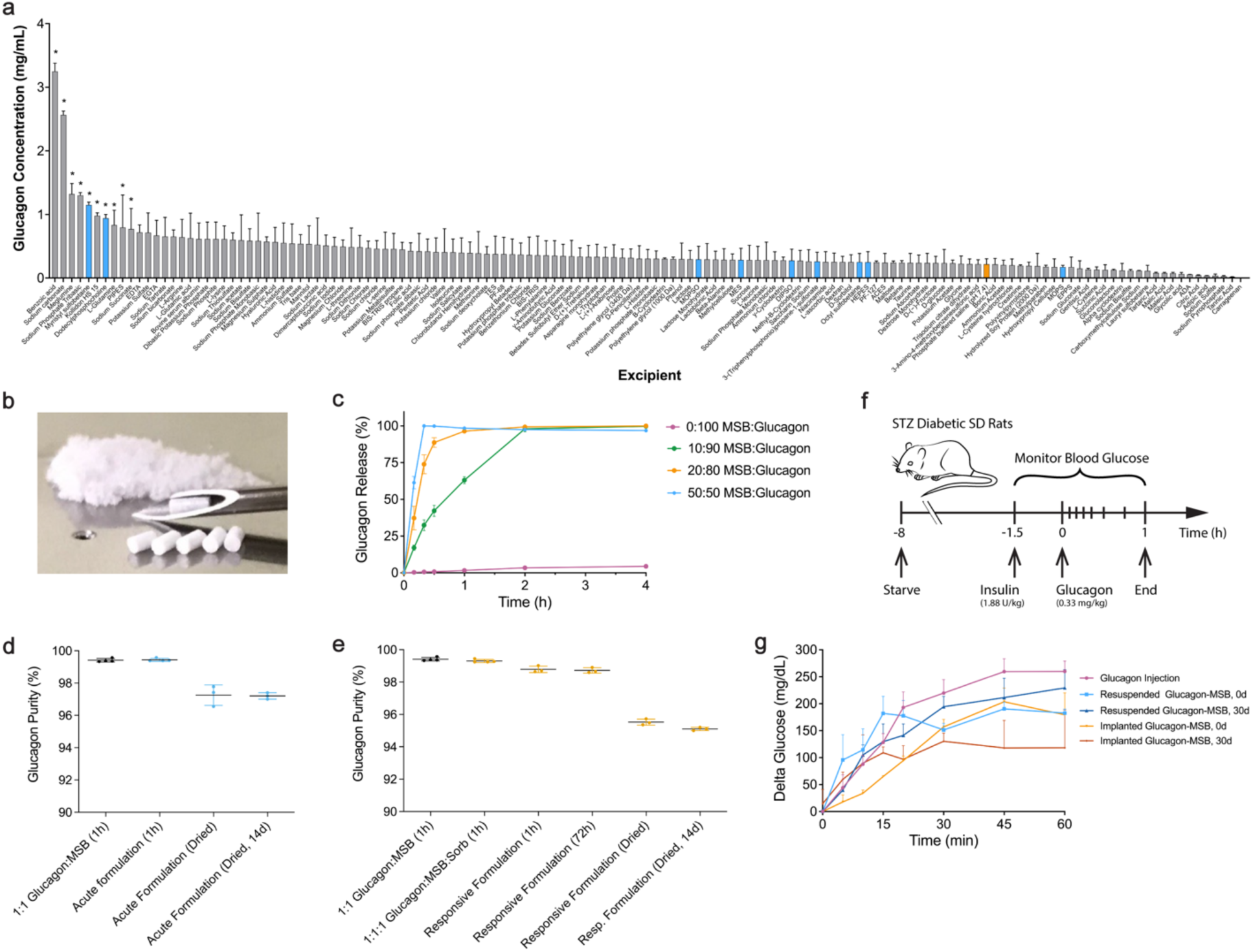
Glucagon formulation. a) In vitro screen to identify solid excipients capable of enhancing glucagon dissolution. Concentration of solubilized glucagon following a 30-minute incubation of lyophilized glucagon in pH 7.4 PBS solutions containing 1 mg/mL of excipient. n ζ 3. * denotes statistically significant differences compared to PBS control denoted in orange. Zwitterionic excipients denoted in blue. b) Photograph of high-density compressed solid glucagon formulations composed of MSB and glucagon. 1 mm diameter tablets loaded within a 16G hypodermic needle. c) Glucagon release kinetics of solid glucagon tablets containing lyophilized glucagon and MSB at various ratios. n ζ 3. d) Stability (i.e., chemical purity) of acute glucagon formulations under various conditions (i.e., freshly prepared, dried, and following storage). n ζ 3. e) Stability (i.e., chemical purity) of responsive glucagon formulations under various conditions (i.e., freshly prepared, dried, and following storage). n ζ 3. f) Scheme for evaluating the bioactivity of glucagon formulations in diabetic rats. g) Hyperglycemic activity of glucagon formulations administered to diabetic rats. n = 3. Data presented as mean ± SD (a-e) or as mean ± SEM (g). Statistically significant differences indicated as * for p < 0.05.

We evaluated the effect of MSB content on the stability of glucagon following storage under accelerated storage conditions (40°C and 75% relative humidity). HPLC analysis of the MSB-glucagon formulations following a 14-day storage period showed no significant loss in glucagon purity (Fig. 2d, 2e). To confirm that the readily soluble thermostable (ReST) solid glucagon formulation, composed of 20% MSB and 80% glucagon, remained biologically active following long-term storage, we resuspended the solid formulations both prepared fresh and after a 30-day storage period at 40°C, in sterile saline and administered them subcutaneously to diabetic rats at a dose of 0.1 mg glucagon (Fig 2f). A comparable therapeutic effect was observed, both in the magnitude of blood glucose elevation and the rate of onset, suggesting no significant loss in biological activity following long-term (30-day) storage (Fig 2g). The resulting pharmacological effects were also similar to freshly reconstituted (unformulated) lyophilized glucagon, indicating no loss in glucagon activity when formulated with MSB. Finally, we administered the ReST solid glucagon formulations to the subcutaneous tissue of the diabetic rats as a solid injectable tablet through a 16-gauge hypodermic needle (i.e., without prior resuspension in a carrier fluid) to evaluate the kinetics of glucagon dissolution and ensuing hyperglycemia. An immediate increase in blood glucose levels was observed, demonstrating the ability of these solid formulations to elicit a rapid therapeutic effect comparable to that of freshly prepared glucagon solutions (Fig. 2g).

### Design of microneedle patches for rapid mini-dose glucagon delivery

Glucagon mini-dosing has been proposed as an effective treatment for several hypoglycemic conditions including in children with impending or mild hypoglycemia^10^, and prevention of exercise-related hypoglycemia^11^. This form of therapy has been shown to be effective in the preceding use cases with low doses of subcutaneous glucagon, typically in the range of 50 µg to 150 µg ^38^. The advantages of glucagon mini-dose therapy include reduced risk of unwanted weight gain that would otherwise occur with frequent rescue carbohydrate treatments^12^, reduced risk of ensuing hyperglycemia following use of glucose tablets to prevent hypoglycemia during exercise, and reduced risk of hypoglycemia unawareness by mitigating repeated episodes of hypoglycemia^9^. However, widespread adoption of glucagon mini-dose treatment in the home setting requires a painless, convenient, and accurate method of glucagon administration which currently does not exist.

Given the short intervention time windows for preventing onset of mild hypoglycemia and progression of mild to severe hypoglycemia in patients with T1D (Table 3), we pursued a microneedle patch design that incorporates the ReST formulation directly into a super-swellable polymer microneedle structure to provide the shortest possible delay between transdermal patch application and glucagon liberation, dissolution, and transport to the bloodstream (Fig. 3a). The patch measures 1 cm by 1 cm in size and features a 14-by-14 array of drug-loaded microneedles, each measuring 1 mm in height (Fig. 3b). Specific levels of glucagon loading within the microneedle tips, ranging from 48.2 µg ± 8.1 µg, 152.3 ug ± 24.7 µg, 299.8 ug ± 51.4 µg, were achieved by adjusting the ratio of drug formulation to polymer matrix (Fig. 3c). The drug loading within the tips is highly reproducible, with variability increasing proportional to the absolute amount of drug loaded. A small ∼2% loss of glucagon purity was observed in the dried microneedle patches, presumably due to dehydration of the solution-casted microneedle patches (Fig. 2c).

**Fig. 3.**
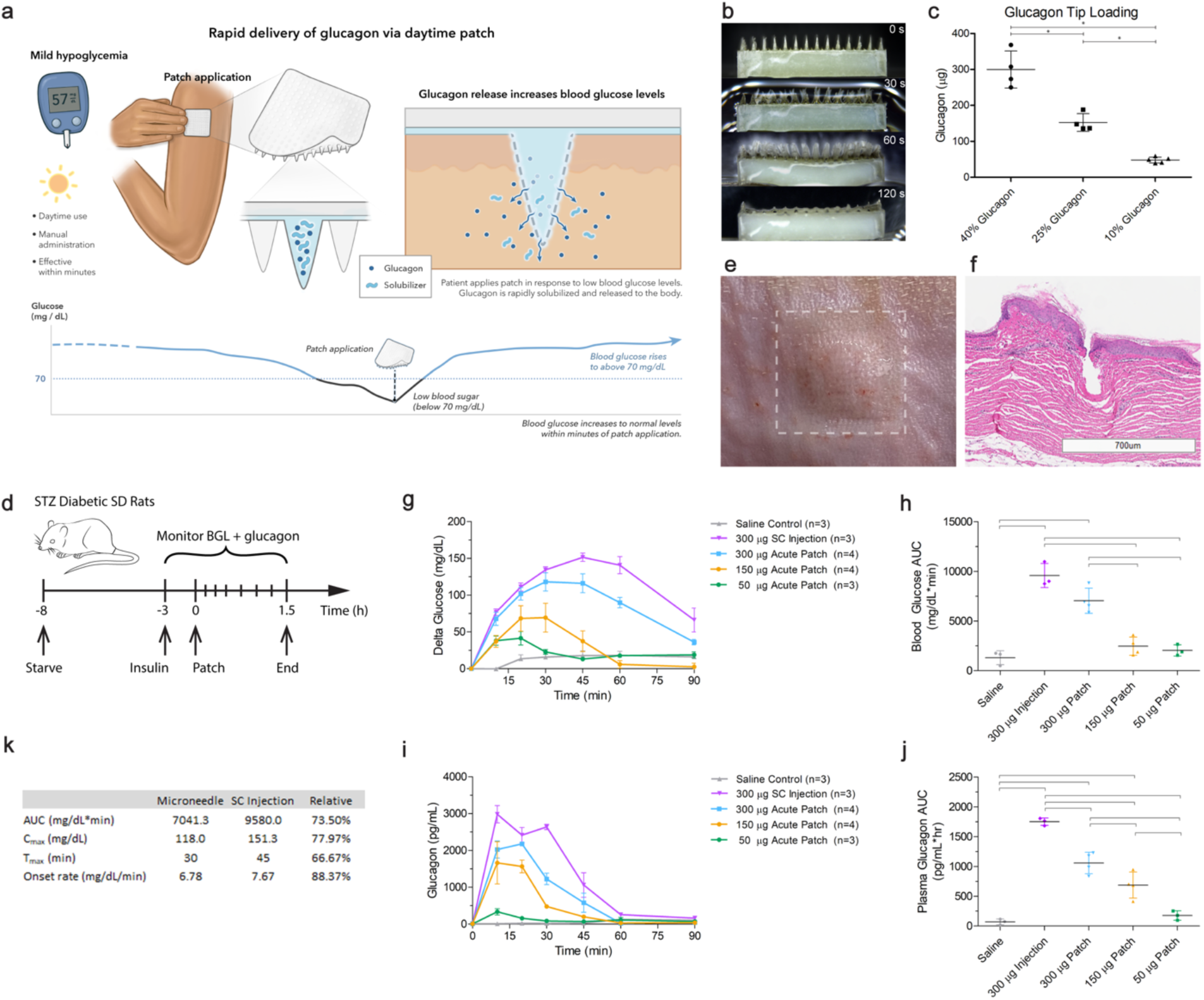
Microneedle patch for glucagon mini dosing. a) concept of glucagon mini-dose delivery using a microneedle patch. b) photograph of microneedle patch before and after immersion in PBS. c) glucagon loading within the microneedle tips. n ζ 4. d) scheme for testing pharmacokinetics and pharmacodynamics in diabetic rats. e) photograph of rat skin following application of microneedle patch. f) H&E-stained tissue section showing microneedle penetration through the stratum corneum of pig skin. g) change in glucose levels following transdermal patch application of microneedle patches loaded with different levels of glucagon, compared to saline control and SC injection of glucagon solution. h) AUC of blood glucose levels following transdermal patch application of microneedle patches loaded with different levels of glucagon, compared to saline control and SC injection of glucagon solution. i) change in plasma glucagon levels following transdermal patch application of microneedle patches loaded with different levels of glucagon, compared to saline control and SC injection of glucagon solution. j) AUC of plasma glucagon levels following transdermal patch application of microneedle patches loaded with different levels of glucagon, compared to saline control and SC injection of glucagon solution. k) AUC, Cmax, Tmax, and rate of blood glucose level change following treatment with microneedle loaded with 300 µg of glucagon or an equivalent dose administered via subcutaneous injection. Data presented as mean ± SD (c) or as mean ± SEM (g-j). Experimental replicates are indicated for each experiment. Statistically significant differences indicated as * for p < 0.05.

Construction of the microneedle structures using super-soluble and biocompatible hyaluronic acid and gelatin polymers imparts sufficient rigidity and strength to the microneedle structure to enable penetration of the stratum corneum of *ex vivo* pig skin while dry (Figure 3f). Upon contact with neutral pH solutions (e.g., pH 7.4 phosphate buffered saline), the xerogel microneedle structures were observed to readily imbibe liquid leading to rapid expansion of microneedle size within 60 seconds, and complete dissolution of both the microneedles and the encapsulated glucagon within 120 seconds (Figure 3b).

### Transdermal delivery and efficacy of glucagon mini-doses in rats

The pharmacokinetics and pharmacodynamics of the glucagon mini-dose microneedle patches was assessed in diabetic rats (Fig. 3d). Transdermal application of the patches loaded with 50 µg, 150 µg, or 300 µg of glucagon resulted in an increase in blood glucose levels detectable within 10 minutes following administration, and a T_max_ of 30-34 minutes (Fig. 3g). The magnitude of the hyperglycemic effect, determined by change in blood glucose levels and area under to blood glucose-time curve (AUC_BG_), was found to increase in a dose-dependent manner (Fig 3h). The hyperglycemic effects observed in the treated rats were further accompanied by a similar dose-dependent increase in plasma glucagon levels (Fig. 3i, 3j). While transdermal delivery of 300 µg of glucagon from the microneedle patch resulted in similar pharmacokinetics and onset of biological activity compared to treatment with an equivalent dose of glucagon administered *via* subcutaneous injection, the total bioavailability and bioactivity of the patches were reduced at 73% and 60%, respectively (Fig. 3k). This reduction is likely due to a combination of incomplete systemic absorption of the drug in the dermal tissue, and a small reduction (∼2%) in drug stability in the microneedle dosage form (Fig. 2c). Critically, the amount of glucagon delivered by the patches and the resulting hyperglycemic effect was found to be highly reproducible.

Transdermal application of the microneedle patches to the skin appears to be well-tolerated, with no signs of redness or permanent damage to the skin surface (Fig. 3e). Histological analysis of the application site revealed successful penetration of the stratum corneum, and no signs of swelling, irritation, or inflammation (Fig. 3f). Taken together, these results suggest that the glucagon mini-dose microneedle patches can pierce the skin of the diabetic rats to access the underlying interstitial fluid leading to accurate, dose-dependent, and rapid systemic delivery of glucagon that is aligned with the timing needs to prevent either onset of mild hypoglycemia (i.e., 44 to 60 minutes) or progression of mild to severe hypoglycemia (i.e., 16 to 25 minutes) in type 1 diabetes patients during the day and at night (Fig. 1, Table 3).

### Design of glucose-responsive microneedle patches for hypoglycemia-triggered glucagon delivery

Automated hypoglycemia-triggered glucagon delivery promises to mitigate the risk of hypoglycemia and eliminate the burden of persistent blood glucose monitoring and manual intervention for patients and their caregivers, particularly during evenings when patients are asleep. Several chemically-driven glucose-responsive systems have recently been reported by our group, and others ^39–41^, however these devices share critical design architectures that may impede clinical translation including (1) use of a non-specific hypoglycemia-triggering mechanism based on phenylboronic acid, which is known to interact with interfering sugars and compounds commonly found in the body (e.g., fructose, galactose, mannose, sialic acid)^42^, and glucagon loading strategies that are prone to continuous drug elution at euglycemic and even hyperglycemic conditions which may exacerbate hyperglycemia and lead to hyperglucagonemia. To overcome these challenges we sought to develop a prophylactic glucose-responsive wearable patch that provides automated and persistent glucagon delivery during episodes of nocturnal hypoglycemia (Fig. 4a). Critically, this system leverages a novel hypoglycemia-responsive mechanism based on the glucose-oxidase enzyme, which is highly specific for glucose, and exhibits no drug release prior to hypoglycemia triggering. Thus, the system can be safely applied prior to sleep, worn throughout the night to protect against nocturnal hypoglycemia, and discarded in the morning.

**Fig. 4.**
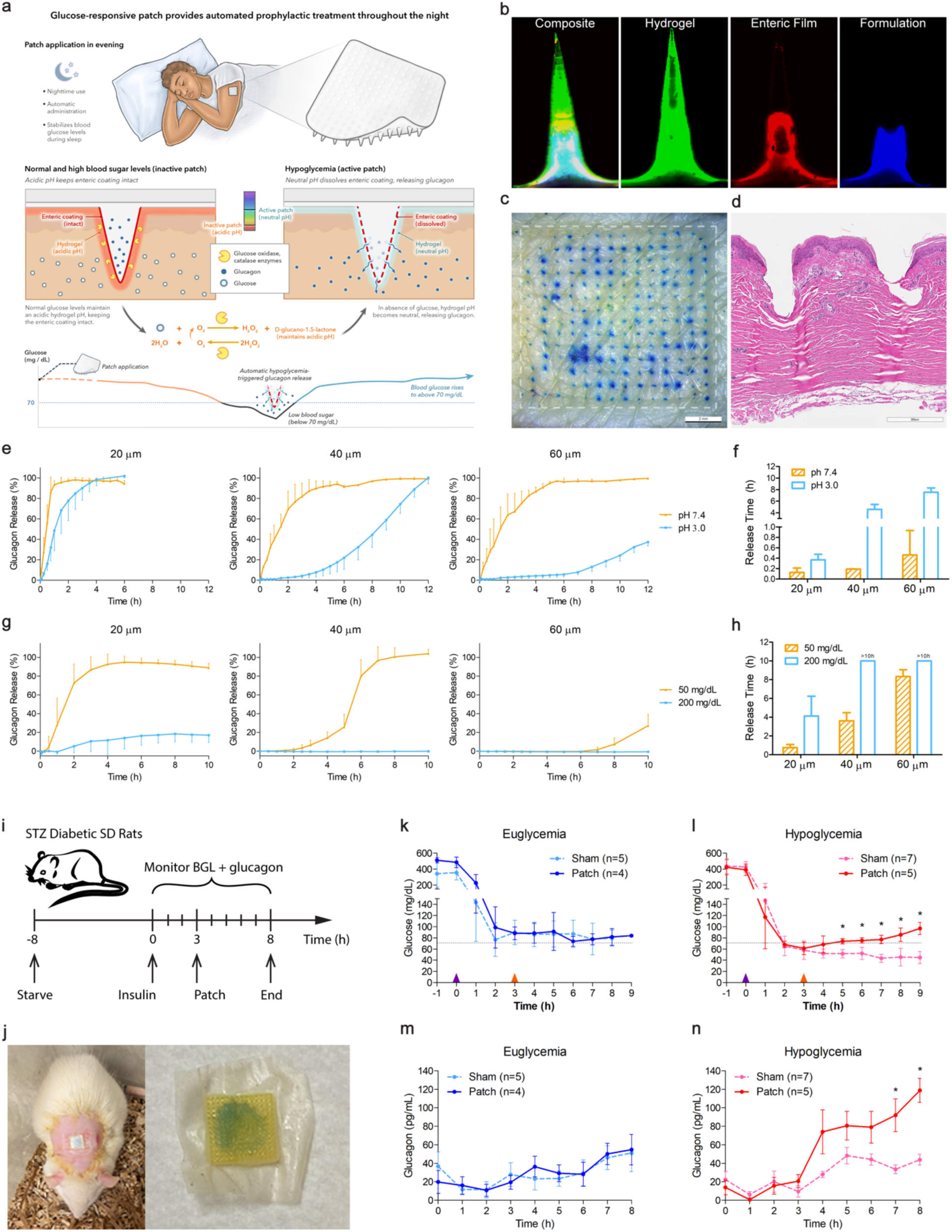
Microneedle patch for hypoglycemia-triggered glucagon delivery. a) Concept of prophylactic treatment and mechanism of hypoglycemia-triggered glucagon release. b) confocal microscopy images of microneedle structure. Enzyme-functionalized hydrogel microneedle structure is shown in green. Enteric film is shown in red. ReST glucagon formulation is shown in blue. c) Photograph of pig skin following application of transdermal microneedle patch. Punctuated blue dots show the penetration of methylene blue dye into the skin following microneedle application. d) H&E-stained tissue section showing microneedle penetration through the stratum corneum of pig skin. e-f) in vitro glucagon release kinetics and release times from the microneedle patch (20, 40, 60 um thick enteric polymer film) under different pH buffers. n ζ 3. g-h) in vitro glucagon release kinetics and release times from the microneedle patch (20, 40, 60 um thick enteric polymer film) under different pH buffers. n ζ 3. i) Scheme for pharmacokinetic and pharmacodynamic evaluation in diabetic rats. j) Photograph of microneedle patch during and after transdermal application. k-l) Blood glucose levels before and after transdermal application of the glucose-responsive glucagon patch. Insulin was administered at 0 h. Microneedle patch was applied at 3 h. m-n) Plasma glucagon levels before and after transdermal application of the glucose-responsive glucagon patch. Insulin was administered at 0 h. Microneedle patch was applied at 3 h. Data presented as mean ± SD (e-h) or as mean ± SEM (k-n). Experimental replicates are indicated for each experiment. Statistically significant differences indicated as * for p < 0.05.

The glucose-responsive microneedle patch is comprised of four distinct layers: (1) A crosslinked hydrogel matrix containing glucose oxidase and catalase enzymes, (2) the ReST solid glucagon formulation, (3) a conformal enteric polymer film that acts as a pH-sensitive barrier between the hydrogel and the glucagon formulation layers, and (4) a solid epoxy backing that provides structural support to the microneedle patch. Confocal fluorescence microscopy of the microneedle patch revealed the distinct hydrogel (green), enteric polymer film (red), and glucagon formulation (blue) layers within the microneedle structure (Fig. 4b). The ability of the microneedles to penetrate the skin was assessed visually following transdermal application to *ex vivo* pig skin followed by topical application of methylene blue solution (Fig. 4c). The visible array of blue dots reveals the location of microneedle penetration through the stratum corneum and localized dye uptake into the pig skin. Histological analysis further revealed microneedle penetration through the stratum corneum, epidermis and into the subdermal layers of the skin (Fig. 4d).

The glucose-responsive mechanism that enables the hypoglycemia-triggered release of glucagon is driven by a pair of enzymatic reactions wherein (1) glucose oxidase catalyzes the oxidation of β-d-glucose into D-glucono-1,5-lactone and hydrogen peroxide, and (2) catalase breaks down the hydrogen peroxide byproduct into water and a regenerated oxygen molecule, which is required for propagation of the enzymatic reactions (Fig. 4a). The final D-glucono-1,5-lactone product is rapidly hydrolyzed into gluconic acid leading to a drop in pH in the local microenvironment. An abundance of glucose during periods of euglycemia and hyperglycemia drives this pair of enzymatic reactions forward resulting in ample production of gluconic acid and maintenance of an acidic microenvironment within the hydrogel matrix. This low pH environment inhibits the dissolution of the enteric polymer film, thereby preventing glucagon release from the microneedle patch. Conversely, the diminished interstitial glucose concentration during periods of hypoglycemia decreases the production of gluconic acid leading to pH neutralization within the hydrogel, rapid dissolution of the enteric polymer film, and release of the glucagon from the microneedle patch. The solid nature of the MSB-glucagon formulation facilitates high drug stability during storage, structural integrity during transdermal application, and timely drug delivery due to its rapid and enhanced solubility in neutral pH solutions.

To confirm the pH-mediated glucose-responsive drug release mechanism, patches constructed using 20 µm, 40 µm and 60 µm-thick shellac enteric polymer films were incubated in buffered acidic or neutral-pH release medium at 37°C under agitation to simulate *in vivo* euglycemic and hypoglycemic conditions, respectively. Drug release kinetics was determined by measurement of glucagon concentrations in the supernatant over a 12-hour period (Fig. 4e). The release time of each patch, quantified as the time required to achieve 10% of total drug release, was determined for both acidic and neutral pH release conditions, and the ratio was used to calculate the device’s responsiveness (R = T_pH 3.0_/T_pH 7.4_) (Fig. 4f). All patches incubated in pH 7.4 release medium (simulated hypoglycemia) exhibited rapid glucagon release in as little as 10 minutes. In contrast, the patches exhibited significantly longer release times (i.e., 0.5, 4, and 7 hours for 20 µm, 40 µm and 60 µm devices, respectively) when incubated in pH 3.0 (simulated euglycemia) conditions; representing a 3-fold, 24-fold to 42-fold longer release in acidic pH conditions compared to neutral pH conditions. These results confirm the proposed pH-sensitive drug release mechanism of this patch system and identify enteric polymer film thickness as a critical parameter in controlling drug release times.

Next, we examine the in vitro glucose-responsive glucagon release kinetics by incubating the patches in pH 7.4 phosphate-buffered saline supplemented with 50 mg/dL or 200 mg/dL glucose at 37°C under agitation to simulate *in vivo* hypoglycemic and euglycemic conditions, respectively. Drug release kinetics was determined by measurement of glucagon concentrations in the supernatant over a 10-hour period (Fig. 4g, 4h). As predicted by the pH-responsive behavior of the system, glucagon release times were faster in low glucose conditions (50 mg/dL) compared to high glucose conditions (200 mg/dL) and were dependent on the enteric polymer chemistry and film thickness. Patches with a 20 µm-thick enteric polymer film exhibited rapid glucagon release within 30 minutes following incubation in low glucose conditions, and gradual glucagon release starting at 2 hours in high glucose conditions. Increasing the enteric polymer film thickness to 40 µm and 60 µm progressively delayed glucagon release times in low glucose conditions to 2 and 4 hours while completely suppressing glucagon release for at least 10 hours. These results suggest that an enteric polymer film thickness between 20 µm and 40 µm may be most suitable to minimize glucagon delivery over the course of the night under euglycemic conditions while providing for an acceptable glucagon release kinetic during hypoglycemic conditions in line with patient needs (Table 3).

### Stability of glucagon formulation in glucose-responsive microneedle patches

The stability of the ReST solid glucagon formulation was evaluated after loading into the glucose-responsive microneedles. Sorbitol was incorporated into the ReST glucagon formulation to increase solution viscosity and facilitate loading into the responsive microneedle structures, and did not lead to a loss in glucagon purity (Fig. 2c). Incorporation of the formulation into the responsive microneedle patches via solution casting resulted in a <1% loss in glucagon purity while wet (1 and 72 hours), followed by a ∼2% loss in drug purity after complete drying. No further change in the chemical purity of the loaded glucagon was observed after a 14-day incubation of the responsive microneedle patches at 40°C and 75% relative humidity.

### Hypoglycemia-triggered glucagon delivery in diabetic rats

Hypoglycemia-triggered glucagon delivery from the microneedle patches was evaluated in diabetic rats (Figure 4i). The rats were fasted for 8 hours and subsequently treated with recombinant human insulin and ultra-long-acting Insulin Degludec^®^ to induce prolonged euglycemia or hypoglycemia. Stable euglycemia or hypoglycemia was achieved 2 hours after insulin treatment and persisted out to 9 hours following administration (Fig. 4k, 4l). Transdermal microneedle patches were applied to the skin 3 hours after insulin administration and secured in place for the duration of the experiment.

Application of the microneedle patches to euglycemic rats had no significant effect on blood glucose levels or plasma glucagon levels compared to euglycemic animals treated with sham devices (Fig. 4k). In this regard, blood glucose levels remained near-constant between 70 mg/dL and 120 mg/dL both before and after transdermal patch application and plasma glucagon levels of both patch and sham-treated euglycemic animals gradually increased from ∼20 mg/dL to ∼50 mg/dL over the course of the experiment (Fig. 4m).

Application of the microneedle patches to hypoglycemic rats induced an immediate and sustained increase in blood glucose levels resulting in a return to euglycemia within 1-2 hours after patch application (Fig. 4l). The kinetics of glucagon release were aligned with the timing needs for automated treatment of nocturnal severe hypoglycemia as reflected in Fig. 1 and Table 3. The blood glucose levels of these animals remained within the euglycemic range for the remainder of the experiment without inducing hyperglycemia (>140 mg/dL). This rise in blood glucose levels following patch application was accompanied by a significant elevation in plasma glucagon levels (Fig. 4n). Blood glucose levels of hypoglycemic animals treated with sham devices remained within the hypoglycemic range and continued to gradually decline for the remainder of the experiment. Plasma glucagon levels in this experimental group were comparable to those observed in the patch and sham-treated euglycemic animals.

### Discussion CGM analysis

Previous studies have confirmed that hypoglycemia is common in patients with T1D and can be prolonged^7,17–19^. However, these findings are insufficient for specifying the engineering design requirements of new hypoglycemic interventions. Here we performed a more detailed analysis of the kinetics of hypoglycemic events to identify a set of engineering design requirements that can help guide the translational development of novel and situation-specific hypoglycemic interventions. We sought to (1) characterize hypoglycemia as distinct events rather than simplified *’time-spent-in hypoglycemia*’ or *’percent nights with hypoglycemia*’ parameterizations, (2) provide holistic treatment of the data by including pre-hypoglycemic periods and associated follow-on hypoglycemic events in the analysis, (3) extract pertinent features for identifying windows of opportunity for glucagon intervention including time-to-treatment, duration and rate of hypoglycemia onset, duration in mild hypoglycemia before severe hypoglycemia, and duration of hypoglycemic events, and (4) expand this analysis to include different levels of hypoglycemia severity, different times of occurrence, and both well-controlled and general T1D patient populations.

Results from our analysis show that a surprisingly large proportion (i.e., 20-30%) of hypoglycemic episodes are follow-on events, suggesting that only 70-80% of all complete events are independent and the total duration of these independent events may be longer when accounting for the associated follow-on events. Roughly 15-20% of parent events were associated with one follow-on event, and parent events with more than one follow-on event occur at a progressively lower frequency. This is likely due to under-treatment of an initial hypoglycemic event leading to failure to maintain euglycemia. Perhaps unsurprisingly, patients with well-controlled diabetes experience ∼14% fewer independent hypoglycemic events compared to the general T1D population, and these instances are associated with ∼10% fewer follow-on events suggesting that patients with well-controlled T1D experience higher success at maintaining euglycemia following an initial hypoglycemic event.

Analysis of hypoglycemic event trajectories in the CGM data shows that the rate and duration of hypoglycemia onset, duration of hypoglycemia, and time to hypoglycemia treatment differ based on event severity (mild vs. severe hypoglycemia), time of hypoglycemia onset (daytime vs. nighttime), and patient population (general vs. well-controlled disease). The rate of glucose decline leading to hypoglycemia onset was found to be faster for daytime events compared to nocturnal events, and in severe cases compared to mild cases. The former may be due to exercise or the use of rapid-acting insulins during the day while the latter may be a direct result of the dose-dependent effect of insulin which affects both the degree and rate of glucose decline. Surprisingly, these differences only translated to small changes in the average pre-hypoglycemia duration, which ranged between 1.0 hours to 1.3 hours in the general T1D population and 44 to 47 minutes in the well-controlled T1D population. This pre-hypoglycemic period represents a largely invariable 44 or 60-minute window of opportunity to prevent impending hypoglycemia following the administration of excessive insulin, either through glucagon mini-dose treatment or other interventions.

In comparison, the average duration of mild hypoglycemia preceding the onset of a severe hypoglycemic event was found to be significantly longer during the night (i.e., ∼16 minutes versus 25 minutes), and represents the second window of opportunity for severe hypoglycemia prevention once mild hypoglycemia has been detected. This period is of particular importance for daytime events where glucagon mini doses may be manually administered shortly after the patient is alerted to low glucose levels to mitigate the risk of a severe event.

The base utility of glucose-responsive or closed-loop glucagon interventions is to provide automated glucagon treatment faster than what is currently achieved using manual intervention. Our findings show that this window of opportunity is highly variable. In general, the average duration of parent hypoglycemic events is longer for severe cases compared to mild cases, during nocturnal events compared to daytime events, and in the general T1D population compared to subjects with well-controlled T1D. These durations can be substantial, ranging from 26 minutes for daytime mild hypoglycemia in subjects with well-controlled disease to 1 hour and 48 minutes for severe nocturnal hypoglycemia in the general T1D patient population.

Finally, the average time to glucagon administration, if detected, ranged from ∼37.5 minutes in patients with well-controlled diabetes to over 50 minutes in the general population (although this difference was not significant for nighttime events). In some instances, multiple glucagon administration events were observed, suggesting that these patients had trouble maintaining euglycemia following a single bolus administration of glucagon. The average time to last glucagon treatment ranged between 53.5 minutes to over 1 hour. This suggests that patients can struggle for prolonged periods of time to control glycemic levels. These observations highlight the clinical utility of automated glucagon system with the ability to provide either multiple cycles or persistent glucagon delivery following hypoglycemia-triggering.

## Formulation

The ReST glucagon formulations identified here overcome the longstanding challenge of developing a storage-stable glucagon formulation that can be administered directly to the body without prior reconstitution. Previous efforts to develop such a formulation focused on stabilizing the glucagon peptide in dilute aqueous and non-aqueous solutions, or low-density powders. Here, we pursued an alternative approach by identifying high-density solid glucagon formulations that rapidly dissolve in neutral pH solutions such as interstitial fluid, to enable storage and administration as a compact dry solid. From a library of 141 water-soluble solid excipients, we identified 10 compounds capable of enhancing glucagon dissolution in pH 7.4 PBS (Fig. 2a, Table S1). From these hits, MSB was identified as a promising excipient based on its ability to enhance solution glucagon concentration by up to 500% compared to unformulated glucagon, and its pH-independent mechanism of action (Fig. S6). A high-density solid formulation composed of 20% MSB and 80% lyophilized glucagon was able to start dissolving within minutes following incubation in a neutral pH buffer.

The solid nature of the ReST glucagon formulation contributes to its inherent stability during storage. We showed that the formulation exhibits minimal chemical and physical degradation following storage at 40°C for up to 14 days and preserved biological effect following 30-day storage. This dry formulation exhibits exceptionally high drug density, up to 80% by weight, which enables space efficient loading into volume-constrained drug delivery devices (e.g., injectable implants, ingestible micro-injectors, or transdermal microneedle patches^28–30^). Administration of high-density ReST pellets into the subcutaneous tissue of diabetic rats elicited rapid peptide dissolution and hyperglycemic effects on timescales that are comparable to subcutaneous injection. Thus, high-density ReST formulations administered by next-generation delivery devices may enable new modes of glucagon therapy so long as the kinetics of formulation administration are in line with the application-specific windows of opportunity identified by the CGM analysis.

### Demonstration 1: Acute Patch

In a first demonstration, we developed painless self-administrable microneedle patches capable of rapid glucagon delivery within the timing needs required to prevent onset of mild hypoglycemia and mitigate progression of mild hypoglycemia to severe hypoglycemia. The high-density and solid nature of the ReST glucagon formulation enabled (1) facile drug loading into the dry polymer microneedles and (2) loading of clinically relevant glucagon doses within a compact 1 cm by 1 cm patch, and remarkable glucagon stability and preserved biologically active even after storage at 40°C for 14 days. Accurate and reproducible control over drug loading within the microneedle structures translated to reproducible dose-dependent drug exposure and hyperglycemic effect in diabetic rats following transdermal patch application. Surprisingly, the solid glucagon formulation delivered by the microneedle patch resulted in a rapid increase in BGL which is comparable to an equivalent dose of liquid glucagon solution injected into the subcutaneous tissue (i.e., significant increase in BGL by 10 minutes and T_max_ by 30 to 45 minutes).

This system addresses several delivery challenges that have thus far impeded widespread adoption of glucagon mini-dose treatment of mild hypoglycemia including the ability to deliver glucagon (1) in a painless and easy-to-use manner from a compact and portable device, (2) at therapeutically relevant and reproducible doses, and (3) with rapid kinetics that are suitable for both prevention of mild hypoglycemia onset in well-controlled and general T1D populations, or progression to severe hypoglycemia when administered at the onset of pre-hypoglycemia or mild hypoglycemia respectively.

### Demonstration 2: Responsive Patch

This system provides automated and persistent glucagon delivery following hypoglycemia-triggering. At the heart of this system is a novel hypoglycemia-responsive drug-release mechanism based on the highly specific glucose oxidase enzyme, which prevents both premature elution of glucagon to the body (even at low levels) during periods of euglycemia or hypoglycemia, and accidental triggering by non-specific analytes. Both challenges limit the clinical translation of previously reported systems^39,40^.

This system further leverages the ReST glucagon formulation to achieve high drug loading and compact patch size. The solid nature of this formulation facilitates easy integration with the microneedle architecture, preserves rigidity of the dry xerogel microneedles, and circumvents the challenge of stabilizing liquid glucagon formulations during long-term storage. The rapid and pH-independent dissolution of this formulation in physiological fluids, including interstitial fluid wicked into the swollen hydrogel microneedles following transdermal application, uniquely enables the timely release of glucagon from the microneedle patch following exposure to low glucose levels.

The glucose-responsive drug release kinetics (or lack thereof) from the patch in hypoglycemic and euglycemic conditions is dependent on the pH-responsive dissolution behavior of the enteric film barrier. Pharmaceutical enteric polymer films are formulated to remain insoluble in acidic pH environments below the material’s pK_a_, and soluble in more neutral pH environments; however, these materials neither dissolve instantaneously at neutral pH nor do they remain indefinitely insoluble at low pH ^43^. Therefore, judicious selection of enteric polymer composition (including pKa and molecular weight) and film thickness is required to achieve the desired glucose-responsive drug delivery profiles for the prophylactic treatment of nocturnal hypoglycemia. Shellac was selected for the construction of the enteric film barrier as this material exhibited the largest pH sensitivity (i.e., the ratio between release time at pH 5.0 and pH 7.4) compared to other enteric polymer systems. By incorporating a 20-micron thick shellac barrier within the microneedle patch, we were able to achieve an average response time of 45.5 minutes, which falls within the window of opportunity to provide automated glucagon delivery following the onset of nocturnal hypoglycemia in T1D patients. Further optimization of this system for clinical translation will be informed by the kinetics of glucagon release in humans.

The present system demonstrates a new glucose-specific enzymatic mechanism for achieving automated glucose responsive glucagon delivery. We demonstrate the ability to prevent glucagon elution prior to hypoglycemia triggering, and suitable glucagon delivery kinetics for the prophylactic treatment of nocturnal hypoglycemic events in patients with type 1 diabetes. Future efforts to identify enteric materials with better pH-sensitive dissolution profiles may lead to faster delivery kinetics towards other applications. Alternative microneedle patch architectures including layer-by-layer designs and encapsulation of enterically-coated solid glucagon microparticles may also be explored to facilitate glucagon rescue from multiple nocturnal hypoglycemic events, however considering the high prevalence (i.e., 20% to 30%) of parent hypoglycemic events with one or more follow-on events, a wearable system that provides persistent elution of glucagon following hypoglycemic triggering may be desirable.

## Conclusion

In summary, a retrospective analysis of CGM data from people with T1D revealed heterogeneous event trajectories that clustered around event severity, time of occurrence, and prior history of hypoglycemia. These observations provide insight into the incidence and kinetics of hypoglycemic events (i.e., onset, persistence, and resolution) of hypoglycemic events, and help define windows of opportunity and engineering requirements for novel glucagon interventions. We further identified high-density solid glucagon formulations that are readily soluble and thermo-stable (ReST) and show two implementations that demonstrate the potential to enable new modes of glucagon therapy (i.e., acute glucagon mini-dose treatment of mild hypoglycemia, and an enzymatic hypoglycemia-triggered prophylactic glucagon treatment of nocturnal hypoglycemia), thereby expanding the clinical role of glucagon beyond the emergency setting.

## Materials and Methods

Dulbecco’s Phosphate-Buffered Saline (PBS) was purchased from Gibco by Life Technologies (Woburn, USA). Recombinant human insulin, glucagon, and Insulin Degludec was obtained from Novo Nordisk A/S (Maalov, Denmark). Steel microneedle masters were purchased from Kjul & Co, ApS (Brøndby, Denmark). Shellac (Protect EN RX Cat # 509521020, Lots 5688996 and 5550391) was gifted and purchased from Sensient Pharmaceutical Technologies (St Louis, MO, USA). Pharmaceutical grade 280 kDa hyaluronic acid was gifted from Bloomage Freda Biopharm Co. (Jinan, China). Dextrose, porcine gelatin (Type A), sorbitol, and streptozotocin were purchased from Sigma Aldrich (Saint Louis, USA).

### Analysis of hypoglycemic events in T1D patient CGM datasets

A retrospective analysis of continuous glucose monitoring (CGM) data from five publicly available clinical trial datasets was performed to characterize the kinetics of hypoglycemia onset, progression, and resolution in patients with type 1 diabetes ^44^. The aggregated dataset is comprised of over 89855 days (approximately 79 days per patient) of CGM traces from 1115 patients that are equally represented in sex (50.9% female; 49.1% male) and highly inclusive across all age groups ranging from 3 to 79 years of age. We divide the dataset by patient enrollment criteria adopted by each of the 5 studies. Data from 909 patients are aggregated from 4 datasets and are considered to be representative of the *general type 1 diabetic population*. The remaining 226 patients enrolled in the study by Aleppo et al. are considered representative of type 1 diabetes patients with *well-controlled diabetes* having no recent history of hypoglycemia or hypoglycemia unawareness.

#### Extracting Hypoglycemic Events

The processing pipeline consists of five stages: (1) preprocessing the raw CGM traces, (2) running the traces through a state machine to extract hypoglycemic events, (3) filtering the events to create a clean dataset, and lastly, (4) analysis and (5) plotting the events. Stages 1-4 were conducted using Python 3.9 ^45^. All statistical tests and plotting of data were completed using RStudio ^46^.

To preprocess the raw data, we generate unique identifiers across datasets from the raw CGM to prevent patient data overlap between studies. We extract individual data into separate files, removing files that do not have BGL data below 70 mg/dL to streamline the analysis and reduce memory consumption.

We run the preprocessed data through a state machine to capture complete hypoglycemic events. Each complete event is represented by a CGM trace comprised of (1) a pre-hypoglycemic period, defined as the period starting at the first local maximum preceding a steady decline in glucose levels into hypoglycemia and concluding when glucose levels drop below the hypoglycemic threshold of 70 mg/dL, and (2) the hypoglycemic period during which blood glucose levels remain below 70 mg/dL. The hypoglycemic period may be further characterized by a period of mild hypoglycemia wherein glucose levels remain between 54 mg/dL and 70 mg/dL, and in some cases, a period of severe hypoglycemia wherein glucose levels drop below 54 mg/dL. A hypoglycemic recovery period is defined by a steady rise in glucose levels from a local minimum back above the 70 mg/dL hypoglycemic threshold to euglycemia. Recovery may be due to the administration of exogenous glucagon, as identified in the CGM trace by a rapid and persistent increase in glucose levels exceeding 20 mg/dL within a 45-minute interval, or for other reasons (e.g., ingestion of carbohydrates, etc.). Analysis parameters extracted from each hypoglycemic event include the time of onset and resolution for pre-hypoglycemia, mild hypoglycemia, and severe hypoglycemia; the initial glucose level at the start of the pre-hypoglycemic period; the minimum glucose level during the event; and the time of glucagon treatment if detected.

CGM traces in which glucose levels do not return to euglycemia (70 mg/dL) due to truncation or gaps in the data are deemed incomplete and are excluded from the analysis (22976 cases; 14270 well-controlled cases; 8706 general population cases). 135,386 distinct complete hypoglycemic events were identified in total. The general population generated 95,953 distinct complete hypoglycemic events, while the well-controlled population generated 62,391.

We categorize these remaining complete events into ‘parent’ events, which can be comprised of individual standalone events or as a series of multiple events. Standalone complete hypoglycemic events or the first event in a series are defined as ‘initial’ events, while hypoglycemic events that closely follow a preceding event within a time interval of <= 1 hour are categorized as ‘follow-on’ events. Each ‘parent’ event was categorized by the time (i.e., daytime vs. nighttime events) and the severity (i.e., mild vs. severe hypoglycemia events) of the minimum glucose level of the ‘initial’ hypoglycemia occurrence. Events in which the onset of the ‘initial’ hypoglycemia occurs between 11 pm and 5:59:59 am are considered night-time events while those with onset times between 6 am and 10:59:59 pm are considered daytime events.

#### Hypoglycemic Event Analysis

Clinical parameters were calculated for each event including the incidence rate, the duration of hypoglycemia, time to glucagon treatment, and the rate of hypoglycemia onset:

- *Average duration* equals the average duration in hypoglycemia across a parent event (including parent and follow-on events).
- *Average pre-hypo duration* includes the “pre-hypoglycemic” phase for only the initial event in a parent event.
- *Average duration to the first glucagon treatment* refers to the time between when a patient first drops into hypoglycemia to the time of first treatment in a parent event.
- *Average duration to last glucagon treatment* refers to the time between when a patient first drops into hypoglycemia to the time of last treatment in a parent event.
- *Average mild duration before first severe hypoglycemia* refers to the time between when a patient drops into hypoglycemia to when they first drop into severe hypoglycemia.
- *Average pre-hypoglycemia to first severe hypoglycemia* refers to the time between a patient’s pre-hypo period to the first time they drop into severe hypoglycemia.
- *The rate of hypoglycemia onset* is defined by the difference in glucose levels between the start time of hypoglycemia and pre-hypoglycemia divided by the time taken to drop into hypoglycemia.
- *Average min BGL* is defined as the minimum blood glucose level across the entire event.

#### Statistical Analysis of Hypoglycemic Events

We investigate the statistical significance of the clinical parameters generated in our analysis between well-controlled and general populations. Additionally, we calculate the statistical significance of the clinical parameters between the individual datasets to capture inter-dataset differences. Clinical parameters did not display normal distributions when plotted. We use Kruskal-Wallis as a non-parametric alternative to one-way ANOVA to compare population mean ranks.

### In silico and high throughput excipient screen

To identify a solid glucagon formulation that is capable of rapid dissolution in physiological fluid, we screened a library of pharmaceutical excipients (Table S1) to identify chemical compounds capable of enhancing the solubility of glucagon in neutral-pH phosphate buffer. The library of compounds included in the screen include 141 solid (T_m_ > 37°C) and water-soluble (≥ 1 mg/mL in water) excipients used in FDA-approved parenteral formulations. We supplemented this library with pharmaceutical polymers, surfactants, buffers, and salts used in non-parental formulations.

Stock solutions of each excipient were prepared in PBS (0.01 M, pH 7.4) at a concentration of 1 mg/mL. These solutions were transferred to 2 mL polypropylene deep well plates and frozen at - 80°C for storage. 4 mg of lyophilized glucagon powder was dispensed into each well of a 0.75 mL polypropylene Micronic plate followed by the addition of 0.5 mL of excipient stock solution. The plates were sealed and placed on an orbital shaker (50 rpm) at 37°C for 30 minutes. The supernatant containing solubilized glucagon was separated from the insoluble drug by centrifugation (4000 g, 5 minutes) and filtration (0.22 µm 96-well cellulose acetate filter plate, 1500g, 5 minutes) prior to analysis. The concentration of solubilized glucagon within the supernatant was quantified using RP-HPLC.

The concentration of glucagon was analyzed using an Agilent 1260 Infinity I HPLC equipped with a UV detector, and an Agilent Poroshell 120 EC-C-18 2.7 um 3.0 x 50 mm column (Agilent 699975-302). Samples were loaded onto the column using a mobile phase consisting of 50% acetonitrile and 50% of 0.1% trifluoroacetic acid. Elution from the column was carried out over a 3-minute period with a flow rate of 0.5 mL/min for 3 minutes starting at 50% acetonitrile and ending at 25% acetonitrile. Glucagon was detected using a UV absorbance of 240 nm, with the primary peak eluting at 2.1 minutes. Glucagon concentration was determining the area under the main glucagon peak.

### Formulation and dissolution kinetics of solid glucagon tablets

Excipients were cryo-milled and sieved to a particle size of 63-120 µm (230-120 mesh) before combining with lyophilized glucagon powder. Solid glucagon formulations were manufactured by tablet compaction of the drug-excipient mixture (1 mm diameter die, 400 N compaction force) using an RD10A Natoli Tablet press (St. Charles, MO, USA). Magnesium stearate, a common lubricant used in tableting, was excluded from this process. The solid formulations were stored at 4°C until use.

The dissolution kinetics of the solid formulations was assessed by incubating the solid glucagon formulations in 2 mL of pH 7.4 PBS, stored at 37°C on an orbital shaker. At predetermined timepoints, 200 µL of supernatant was removed for HPLC analysis and an equivalent volume of release medium was replaced. The samples were diluted with equal parts of acetonitrile and analyzed using RP-HPLC.

### Evaluation of glucagon formulation stability

The stability of glucagon formulations was evaluated by packaging the formulations within desiccated and sealed glass vials, followed by incubation at 40°C and 75% relative humidity for 14 to 30 days. An identical batch of formulations was stored at 4°C. The chemical stability of the formulations following incubation was assessed using by measuring glucagon purity using RP HPLC, glucagon structure using circular dichroism, and bioactivity in diabetic rats.

Samples preparation for the chemical purity analysis proceeded by diluting glucagon formulations to a final drug concentration of 250 µg/mL in equimolar acetonitrile and water, and further supplemented with 2 µL of 1M HCl. The chemical purity analysis was performed using the HPLC method described in the “ In silico and high throughput excipient screen” section. Glucagon purity was determined as the percent area of the main peak relative to the total area of all peaks within the RP-HPLC chromatogram

Sample preparation for circular dichroism analysis proceeded by diluting samples to a final concentration of 50 μM in 0.154 mM pH 7.4 PBS. Samples were analyzed using a JASCO circular dichroism using a 2 mm pathlength quartz cuvette.

### Fabrication of acute microneedles

PDMS microneedle molds were prepared by casting a steel microneedle master within a well-mixed solution of PDMS (1:10 ratio of crosslinker to base). The PDMS molds were heated at 60°C for 12 hours followed by the removal of the steel microneedle master.

A polymer stock solution was prepared by dissolving 3 mg of hyaluronic acid and 3 mg of gelatin in 80 μL of 0.35 M HCl by incubating at 37°C for 4 hours. A separate glucagon solution was prepared by dissolving 1, 3, or 6 mg (10%, 25%, and 40% drug loading respectively) of lyophilized human recombinant glucagon and 3 mg of DMPS in 40 μL of 0.35 M HCl. The polymer and glucagon stock solutions were then combined, thoroughly mixed, and centrifuged to remove air bubbles. The mixture was then dispensed into a PDMS microneedle mold and centrifuged at 4000 rpm for 3 minutes. The microneedles were left at room temperature to dry overnight. 40 μL of medical-grade epoxy (Loctite M-21HP) was introduced to the PDMS microneedle mold and centrifuged at 500 rpm for 1 minute. The epoxy was allowed to solidify overnight. The demolded microneedles were attached to a Tegaderm adhesive bandage and stored in a desiccated container at 4°C until use.

### Synthesis of gelatin methacrylate

20 g of gelatin (porcine, type A) was dissolved in 20 mL of 60°C DI water. 10 mL of methacrylic anhydride was added dropwise to the stirred solution. The reaction was allowed to proceed for 2 hours at 60°C. The resulting product was cooled to room temperature and dialyzed against DI water for 3 days at 4°C.

### Synthesis of methacrylated hyaluronic acid

2 g of HA was dissolved in 100 mL of DI water and stirred at 500 rpm for 2 hours at room temperature. 1.6 mL of methacrylic anhydride was added dropwise to the solution. 5 N NaOH was added dropwise to adjust the final pH of the solution to pH 8.5. The solution was protected from light and stirred at 4oC for 24 hours. 2.92 g of NaCl was added to the solution followed by precipitation of the polymer in ethanol. The resulting product was then isolated by centrifugation and washed in ethanol. The product was then redissolved in DI water and dialyzed against DI water for 3 days at 4°C.

### Fabrication of glucose-responsive microneedles

A stock photo-initiator solution was prepared by dissolving 60 mg of Irgacure I2959 in 3.6 mL of DI water and heated at 65°C for 12 hours while protected from light. A polymer stock solution was prepared by dissolving 100 mg of MeHA, 100 mg of GelMA, 4 mg of MBA in 1.6 mL of pH 7.4 PBS. This solution was heated at 37°C for 8 hours to ensure complete polymer dissolution. To this stock, 40 mg of glucose oxidase, 13.3 mg of catalase, and 2.4 mL of photo-initiator stock were added, vortexed and incubated at 37°C for 15 minutes. 4 μL of 25% glutaraldehyde solution was added to the solution and vortexed. The homogenous solution was then centrifuged at 1000 rpm for 3 minutes to remove air bubbles.

80 μL of polymer stock solution was dispensed into each microneedle mold and centrifuged at 4000 rpm for 3 minutes. The molds were protected from light and allowed to dry overnight at room temperature. The dried polymer structure was subsequently exposed to UV irradiation in a UV oven for 15 minutes to facilitate UV-initiated hydrogel crosslinking. Preparation of devices for confocal imaging included 4 mg of 3-5 kDa FITC dextran within the formulation.

A 5% w/w shellac stock solution was prepared by dissolving 60 g Protect ENRX in 276 g of isopropyl alcohol. This solution was applied conformally to the back of the crosslinked hydrogel by spray coating. The enteric film was protected from light and allowed to dry overnight at room temperature. 0.5 mg of Texas red NHS ester was incorporated into the shellac film for devices used for confocal imaging.

A glucagon stock solution was prepared by dissolving 10 mg MSB, 10 mg glucagon, 25 mg sorbitol, and 5 mg FITC-dextran in 40 μL of 0.35 M HCl. 7.6 μL of stock solution was dispensed into each mold and allowed to dry at room temperature overnight in the dark. For devices used in confocal imaging, FITC-dextran was replaced with an equivalent mass of sorbitol and 0.5 mg of Cy5.5-labeled glucagon was incorporated into the formulation for each device.

Finally, 40 μL of medical-grade epoxy (Loctite M-21HP) was introduced to the PDMS microneedle mold, centrifuged at 500 rpm for 1 minute, and allowed to cure overnight at room temperature.

### In vivo evaluation of glucagon formulations in diabetic rats

All animal experiments were approved by and performed in accordance with the Committee on Animal Care at MIT. 200-225 g Sprague Dawley (SAS SD strain 400) instrumented with jugular vein catheters and Instech buttons were purchased from Charles River. The animals were starved for 8 hours and subsequently treated with streptozotocin (STZ) solution, freshly prepared in pH 4.5 50 mM citrate buffer, at a 65 mg/kg dose and administered by intraperitoneal injection.

A 1 U/mL stock solution of human recombinant insulin was prepared by dissolving 1.36 mg of human recombinant insulin in 23.0 mL of pH 8.0 sterile saline. A 1 U/mL stock solution of insulin Degludec was prepared by diluting 100 μl of commercial U100 (100 U/mL) Degludec in 9900 μl of sterile saline. Animals were fasted for 8 hours before treatment with a total insulin dose of 2-6 U/kg, made up of 30% human recombinant insulin and 70% insulin Degludec, to induce stable euglycemia or hypoglycemia. Insulin formulations were administered in separate subcutaneous injections away from the site of patch application. Stable euglycemia or hypoglycemia was achieved 3-hours following insulin dosing. Patches were applied to the shaved and de-pilfered skin of the animals and secured using a Tegaderm bandage. Acute patches were removed 5 minutes after application while responsive patches were left in place for the duration of the experiment. At various timepoints, blood was sampled from the jugular catheter, measured for blood glucose levels using a handheld glucometer, and collected in BD microtainer tubes with K_2_EDTA and 14 μL of 7 TIU/mL (91 kIU/mL, lot SLBT2024) aprotinin. Samples were stored on ice prior to processing. Each sampled was centrifuged at 10,000 rpm for 5 minutes and the plasma was collected and stored at -80°C prior to analysis. Plasma glucagon and c-peptide levels were measured using commercial ELISA kits (Invitrogen) according to the manufacturer’s protocols.

### Statistical Analysis

Unpaired two-sided Student’s t-tests and one-way analysis of variance (ANOVA) with Tukey’s multiple comparisons tests were performed using GraphPad Prism (Version 9.4.1) or Microsoft Excel. A value of P<0.05 was considered statistically significant. Figure captions and text describe the number of replicates used in each study. Figure captions define the center line, and error bars presented in the plots.

## Acknowledgments

We thank the MIT Koch Institute Swanson Biotechnology Center histology and high throughput cores for technical support. Pharmaceutical grade 280 kDa hyaluronic acid was gifted from Bloomage Freda Biopharm Co. (Jinan, China). Concept illustrations by Virginia E. Fulford, Alar Illustrations.

## Author contributions

J.L., G.T. developed general approach and concept. C.B., J.L., developed the hypoglycemia CGM data workflow and analysis. J.L. designed the device. J.L., J.Y.L., S.L., A.M., S.L., S.L, A.L., P.K., J.W.C., developed and characterized the stabilized formulations and the devices. J.L., J.Y.L, S.L., M.K.L., A.M., S.L., A.H., A.V. performed in vivo experiments. J.L., J.Y.L., S.L., M.K.L., A.M., S.L., A.L., P.K., J.F., and A.V. performed analysis on in vitro and in vivo data. J.L., J.Y.L., and the Swanson Biotechnology Center performed histology. U.R., S.T.B, R.L., G.T. had oversight and leadership responsibility for the research activity planning and execution. J.L., C.B., J.Y.L., R.L. and G.T. wrote the manuscript. All authors discussed results and commented on the manuscript.

## Data and materials availability

All data are available in the main text or the supplementary materials. The data used to generate the figures and numerical statistics in this paper can be found as attached or at github and in the Supplementary Information.

## References

1. Workgroup on Hypoglycemia, American Diabetes Association. Defining and reporting hypoglycemia in diabetes: a report from the American Diabetes Association Workgroup on Hypoglycemia. Diabetes Care 28, 1245–1249 (2005).

2. American Diabetes Association. 6. Glycemic Targets: Standards of Medical Care in Diabetes-2021. Diabetes Care 44, S73–S84 (2021).

3. The Effect of Intensive Treatment of Diabetes on the Development and Progression of Long-Term Complications in Insulin-Dependent Diabetes Mellitus. N. Engl. J. Med. 329, 977–986 (1993).

4. Tanenberg, R. J., Newton, C. A. & Drake, A. J. Confirmation of Hypoglycemia in the “Dead-In-Bed” Syndrome, as Captured by a Retrospective Continuous Glucose Monitoring System. Endocr. Pract. 16, 244–248 (2010).

5. Hwang, J. J. et al. Hypoglycemia unawareness in type 1 diabetes suppresses brain responses to hypoglycemia. J. Clin. Invest. 128, 1485–1495 (2018).

6. Seaquist, E. R. et al. Hypoglycemia and diabetes: a report of a workgroup of the American Diabetes Association and the Endocrine Society. Diabetes Care 36, 1384–1395 (2013).

7. Lin, Y. K. et al. Associations Between the Time in Hypoglycemia and Hypoglycemia Awareness Status in Type 1 Diabetes Patients Using Continuous Glucose Monitoring Systems. Diabetes Technol. Ther. 22, 787–793 (2020).

8. Pieber, T. R. et al. DEVOTE 3: temporal relationships between severe hypoglycaemia, cardiovascular outcomes and mortality. Diabetologia 61, 58–65 (2018).

9. Castle, J. R. Is Mini-Dose Glucagon the Answer to Preventing Exercise-Related Dysglycemia? Diabetes Care 41, 1842–1843 (2018).

10. Haymond, M. W. & Schreiner, B. Mini-dose glucagon rescue for hypoglycemia in children with type 1 diabetes. Diabetes Care 24, 643–645 (2001).

11. Rickels, M. R. et al. Mini-Dose Glucagon as a Novel Approach to Prevent Exercise-Induced Hypoglycemia in Type 1 Diabetes. Diabetes Care 41, 1909–1916 (2018).

12. Tetzschner, R., Ranjan, A. G., Schmidt, S. & Nørgaard, K. Preference for Subcutaneously Administered Low-Dose Glucagon Versus Orally Administered Glucose for Treatment of Mild Hypoglycemia: A Prospective Survey Study. Diabetes Ther. 10, 2107–2113 (2019).

13. Pontiroli, A. E. Intranasal glucagon: a promising approach for treatment of severe hypoglycemia. J. Diabetes Sci. Technol. 9, 38–43 (2015).

14. Kedia, N. Treatment of severe diabetic hypoglycemia with glucagon: an underutilized therapeutic approach. Diabetes Metab. Syndr. Obes. Targets Ther. 4, 337–346 (2011).

15. Wilson, L. M. & Castle, J. R. Stable Liquid Glucagon: Beyond Emergency Hypoglycemia Rescue. J. Diabetes Sci. Technol. 12, 847–853 (2018).

16. Weinstock, R. S. et al. Risk Factors Associated With Severe Hypoglycemia in Older Adults With Type 1 Diabetes. Diabetes Care 39, 603–610 (2015).

17. Buckingham, B. et al. Duration of Nocturnal Hypoglycemia Before Seizures. Diabetes Care 31, 2110–2112 (2008).

18. Carlson, A. L. et al. Hypoglycemia and Glycemic Control in Older Adults With Type 1 Diabetes: Baseline Results From the WISDM Study. J. Diabetes Sci. Technol. 15, 582–592 (2021).

19. Juvenile Diabetes Research Foundation Continuous Glucose Monitoring Study Group. Prolonged nocturnal hypoglycemia is common during 12 months of continuous glucose monitoring in children and adults with type 1 diabetes. Diabetes Care 33, 1004–1008 (2010).

20. Müller, T. D., Finan, B., Clemmensen, C., DiMarchi, R. D. & Tschöp, M. H. The New Biology and Pharmacology of Glucagon. Physiol. Rev. 97, 721–766 (2017).

21. Pieber, T. R. et al. Dasiglucagon—A Next-Generation Glucagon Analog for Rapid and Effective Treatment of Severe Hypoglycemia: Results of Phase 3 Randomized Double-Blind Clinical Trial. Diabetes Care 44, 1361–1367 (2021).

22. Newswanger, B. et al. Development of a Highly Stable, Nonaqueous Glucagon Formulation for Delivery via Infusion Pump Systems. J. Diabetes Sci. Technol. 9, 24–33 (2015).

23. Yale, J.-F. et al. Faster Use and Fewer Failures with Needle-Free Nasal Glucagon Versus Injectable Glucagon in Severe Hypoglycemia Rescue: A Simulation Study. Diabetes Technol. Ther. 19, 423–432 (2017).

24. Clinical Evidence. Clinical Review Report: Glucagon Nasal Powder (Baqsimi): (Eli Lilly Canada Inc): Indication: For the treatment of severe hypoglycemic reactions which may occur in the management of insulin treated patients with diabetes mellitus, when impaired consciousness precludes oral carbohydrates [Internet] (Canadian Agency for Drugs and Technologies in Health, 2020).

25. Sherman, J. J. & Lariccia, J. L. Glucagon Therapy: A Comparison of Current and Novel Treatments. Diabetes Spectr. 33, 347–351 (2020).

26. Rickels, M. R. et al. Intranasal Glucagon for Treatment of Insulin-Induced Hypoglycemia in Adults With Type 1 Diabetes: A Randomized Crossover Noninferiority Study. Diabetes Care 39, 264–270 (2016).

27. Pontiroli, A. E., Rizzo, M. & Tagliabue, E. Use of glucagon in severe hypoglycemia is scarce in most countries, and has not been expanded by new ready-to-use glucagons. Diabetol. Metab. Syndr. 14, 193 (2022).

28. Chen, W. et al. Dynamic omnidirectional adhesive microneedle system for oral macromolecular drug delivery. Sci. Adv. 8, eabk1792 (2022).

29. Abramson, A. et al. An ingestible self-orienting system for oral delivery of macromolecules. Science 363, 611–615 (2019).

30. Caffarel-Salvador, E. et al. A microneedle platform for buccal macromolecule delivery. Sci. Adv. 7, eabe2620 (2021).

31. The Diabetes Research in Children Network (DirecNet) Study Group. A Randomized Multicenter Trial Comparing the GlucoWatch Biographer With Standard Glucose Monitoring in Children With Type 1 Diabetes. Diabetes Care 28, 1101–1106 (2005).

32. Aleppo, G. et al. REPLACE-BG: A Randomized Trial Comparing Continuous Glucose Monitoring With and Without Routine Blood Glucose Monitoring in Adults With Well-Controlled Type 1 Diabetes. Diabetes Care 40, 538–545 (2017).

33. Diabetes Research in Children Network (DirecNet) Study Group et al. Continuous glucose monitoring in children with type 1 diabetes. J. Pediatr. 151, 388–393, 393.e1–2 (2007).

34. The Effect of Continuous Glucose Monitoring in Well-Controlled Type 1 Diabetes. Diabetes Care 32, 1378–1383 (2009).

35. Juvenile Diabetes Research Foundation Continuous Glucose Monitoring Study Group et al. Continuous glucose monitoring and intensive treatment of type 1 diabetes. N. Engl. J. Med. 359, 1464–1476 (2008).

36. Wiltshire, E. J., Newton, K. & McTavish, L. Unrecognised hypoglycaemia in children and adolescents with type 1 diabetes using the continuous glucose monitoring system: Prevalence and contributors. J. Paediatr. Child Health 42, 758–763 (2006).

37. Pratley, R. E. et al. Effect of Continuous Glucose Monitoring on Hypoglycemia in Older Adults With Type 1 Diabetes: A Randomized Clinical Trial. JAMA 323, 2397–2406 (2020).

38. El Youssef, J., et al. Quantification of the Glycemic Response to Microdoses of Subcutaneous Glucagon at Varying Insulin Levels. Diabetes Care 37, 3054–3060 (2014).

39. GhavamiNejad, A. et al. Glucose-Responsive Composite Microneedle Patch for Hypoglycemia-Triggered Delivery of Native Glucagon. Adv. Mater. Deerfield Beach Fla 31, e1901051 (2019).

40. Wang, Z. et al. Dual self-regulated delivery of insulin and glucagon by a hybrid patch. Proc. Natl. Acad. Sci. 117, 29512–29517 (2020).

41. Glucose-responsive microneedle patch for closed-loop dual-hormone delivery in mice and pigs | Science Advances. https://www.science.org/doi/full/10.1126/sciadv.add3197.

42. Wu, X. et al. Selective sensing of saccharides using simple boronic acids and their aggregates. Chem. Soc. Rev. 42, 8032–8048 (2013).

43. Liu, F. et al. A novel concept in enteric coating: A double-coating system providing rapid drug release in the proximal small intestine. J. Controlled Release 133, 119–124 (2009).

44. Martin, M. et al. irinagain/Awesome-CGM: Updated release with simulators and enhanced processing. (2021) doi:10.5281/zenodo.4723654.

45. Van Rossum, G. & Drake, F. L. Python 3 Reference Manual. (CreateSpace, 2009).

46. RStudio Team (2020). RStudio: Integrated Development for R. PBC Boston MA.

