## Supplemental Information for "Kinetics of Hypoglycemia in Diabetes Patients Informs Development of New Modes of Glucagon Therapy"

#### Affiliations

**Supplementary Figures**

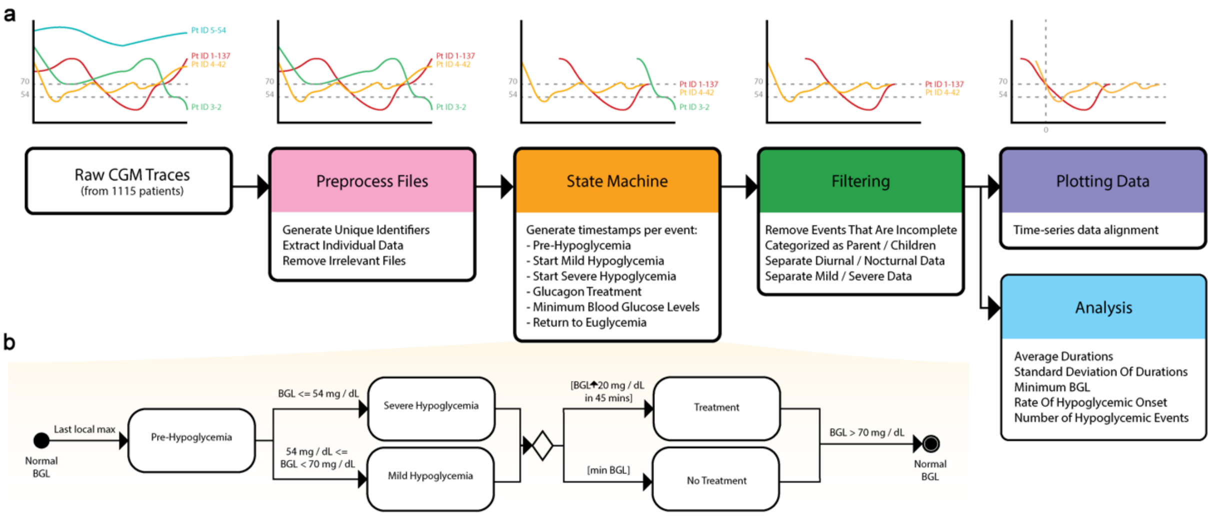


**Fig. S1.** Process workflow of CGM data. a) data workflow and b) state machine diagram.

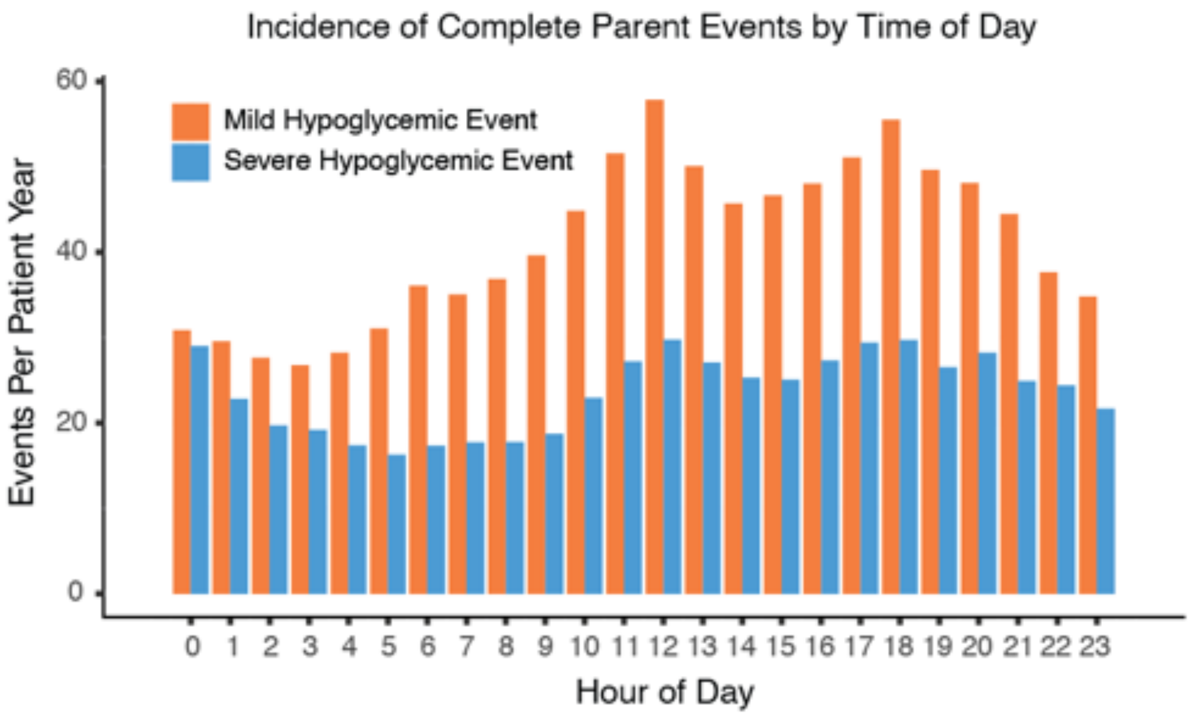


**Fig. S2.** Incidence of complete parent events by time of occurrence. Mild hypoglycemic events in orange. Severe hypoglycemic events in blue.

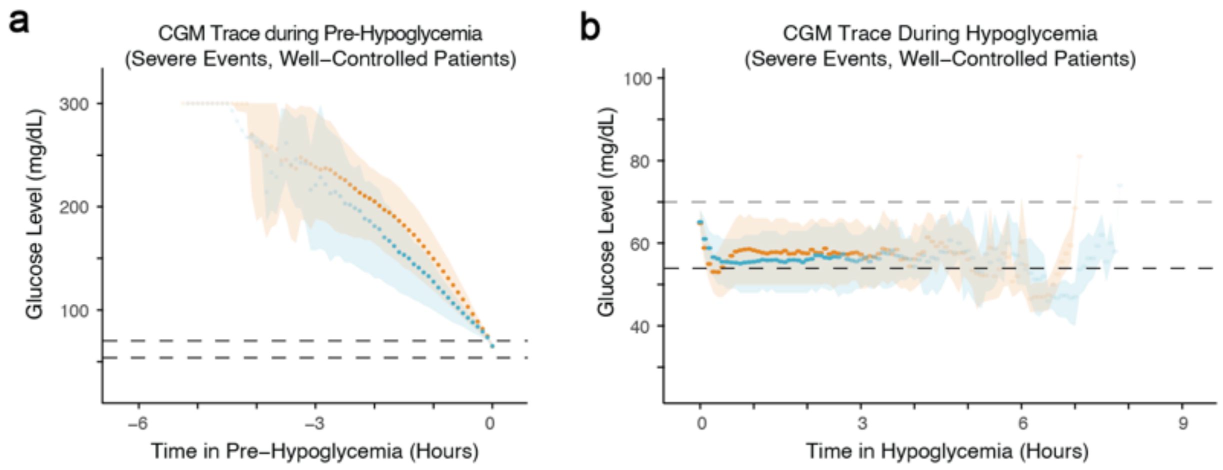


**Fig. S3.** CGM time traces. a) Average (dots, 25th and 75th percentile (shaded areas) of daytime (orange) and nighttime (blue) CGM traces during pre-hypoglycemia. b) Average (dots), 25th and 75th percentile (shaded areas) of daytime (orange) and nighttime (blue) CGM traces during hypoglycemia.

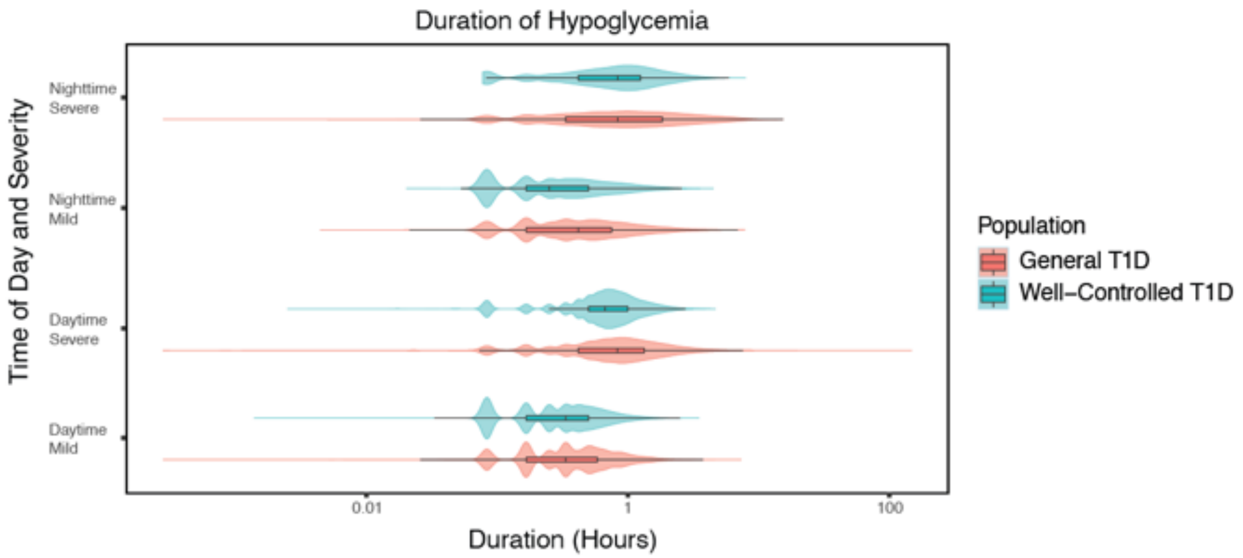


**Fig. S4.** Violin plot of the hypoglycemic event duration in patients with general and well-controlled T1D.

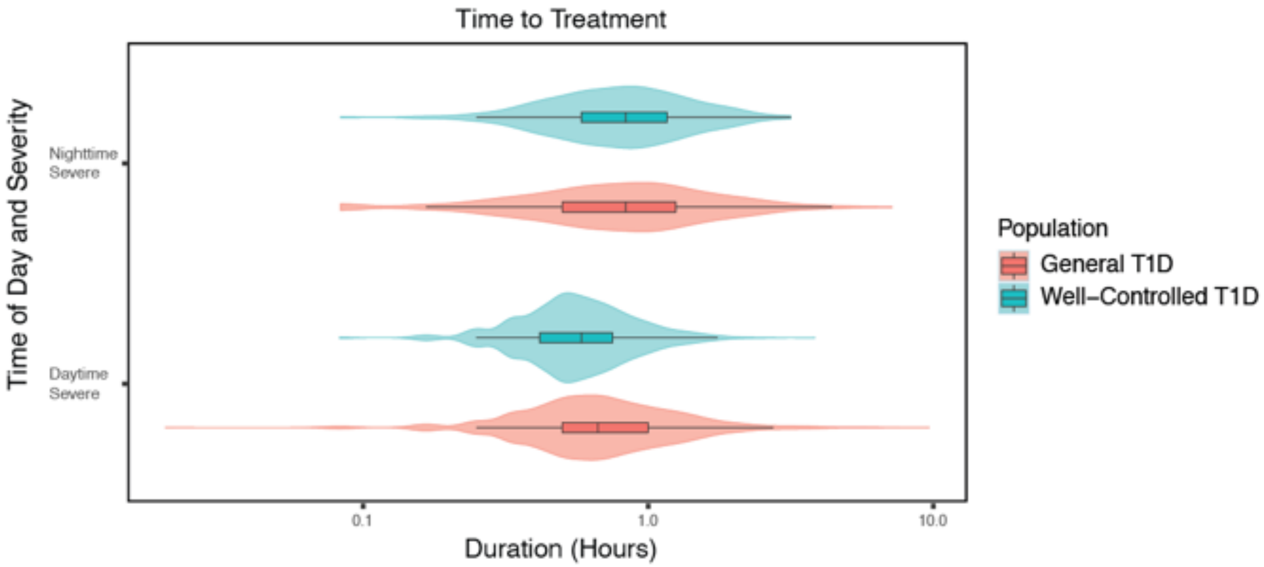


**Fig. S5.** Violin plot of the time to treatment for in patients with general and well-controlled T1D.

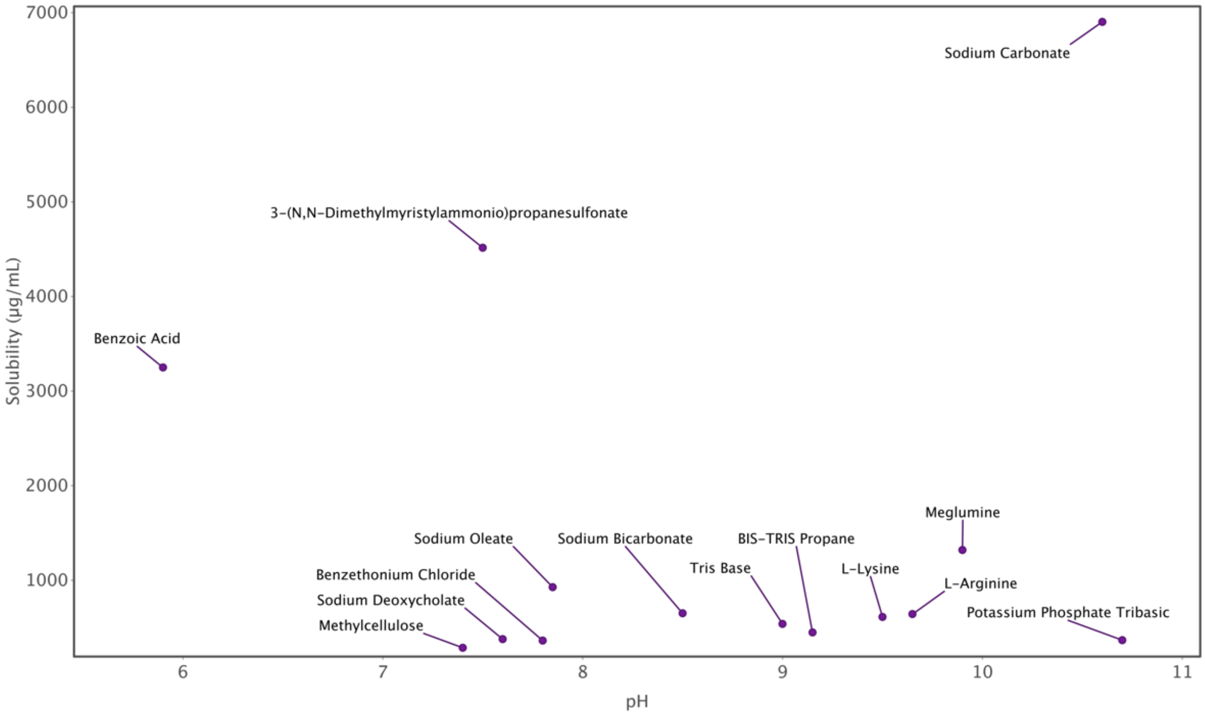


**Fig. S5.** Glucagon solubility versus pH of excipient solutions. Solutions were prepared at a concentration of 1 mg/mL in water.

### Supplementary Tables

#### Table S1.

High throughput screening of a library of water-soluble solid excipients.

| | Excipient | Glucagon concentration | | Fold change vs. PBS | |
|---|---|---|---|---|---|
| | | Avg. (mg/mL) | SD | Avg. (fold) | SD |
| 1 | Benzoic acid | 3.25 | 0.13 | 15.09 | 0.60 |
| 2 | Sodium carbonate | 2.56 | 0.06 | 11.91 | 0.29 |
| 3 | Meglumine | 1.32 | 0.17 | 6.13 | 0.77 |
| 4 | Sodium Phosphate Tribasic | 1.30 | 0.04 | 6.04 | 0.20 |
| 5 | Myristyl sulfobetaine | 1.15 | 0.04 | 5.33 | 0.21 |
| 6 | Kollidon HS 15 | 0.98 | 0.05 | 4.55 | 0.22 |
| 7 | Dodecylphosphocholine | 0.94 | 0.06 | 4.37 | 0.28 |
| 8 | L-Glutamine | 0.83 | 0.23 | 3.87 | 1.08 |
| 9 | PIPES | 0.80 | 0.51 | 3.70 | 2.37 |
| 10 | Sodium Succinate | 0.77 | 0.32 | 3.58 | 1.50 |
| 11 | EDTA | 0.72 | 0.12 | 3.34 | 0.55 |
| 12 | Potassium Sulfate | 0.71 | 0.31 | 3.32 | 1.44 |
| 13 | EGTA | 0.67 | 0.24 | 3.11 | 1.11 |
| 14 | Sodium Tartrate | 0.65 | 0.30 | 3.04 | 1.39 |
| 15 | Sodium bicarbonate | 0.65 | 0.11 | 3.03 | 0.49 |
| 16 | L-Arginine | 0.64 | 0.28 | 2.98 | 1.30 |
| 17 | L-Glutamic acid | 0.63 | 0.39 | 2.91 | 1.82 |
| 18 | Bovine serum albumin | 0.62 | 0.25 | 2.86 | 1.16 |
| 19 | Dibasic Potassium Phosphate | 0.61 | 0.27 | 2.85 | 1.24 |
| 20 | Sodium Phosphite | 0.61 | 0.27 | 2.85 | 1.24 |
| 21 | L-lysine | 0.61 | 0.21 | 2.84 | 0.97 |
| 22 | Sodium Thiosulfate | 0.60 | 0.11 | 2.80 | 0.51 |
| 23 | Sodium acetate | 0.59 | 0.40 | 2.76 | 1.86 |
| 24 | Sodium Bisulfate | 0.59 | 0.19 | 2.72 | 0.88 |
| 25 | Sodium Phosphate Monobasic | 0.58 | 0.44 | 2.72 | 2.03 |
| 26 | Magnesium Sulphate | 0.57 | 0.08 | 2.66 | 0.36 |
| 27 | Hyaluronic Acid | 0.57 | 0.23 | 2.63 | 1.07 |
| 28 | L-histidine | 0.55 | 0.31 | 2.57 | 1.45 |
| 29 | Ammonium Sulfate | 0.54 | 0.31 | 2.52 | 1.44 |
| 30 | Tris base | 0.54 | 0.14 | 2.50 | 0.67 |
| 31 | Mannitol | 0.54 | 0.29 | 2.49 | 1.33 |
| 32 | Sodium Lactate | 0.52 | 0.42 | 2.44 | 1.95 |
| 33 | Dimercaptosuccinic acid | 0.51 | 0.17 | 2.38 | 0.77 |
| 34 | Sodium Iodide | 0.50 | 0.14 | 2.33 | 0.65 |
| 35 | Magnesium Chloride | 0.50 | 0.10 | 2.30 | 0.48 |
| 36 | L-threonine | 0.49 | 0.30 | 2.26 | 1.39 |
| 37 | Sodium Dithionite | 0.49 | 0.19 | 2.25 | 0.86 |
| 38 | Sodium Chlorate | 0.47 | 0.11 | 2.19 | 0.51 |
| 39 | Sodium chloride | 0.46 | 0.13 | 2.14 | 0.59 |
| 40 | L-serine | 0.46 | 0.26 | 2.13 | 1.21 |
| 41 | Potassium Metabisulfite | 0.46 | 0.24 | 2.13 | 1.13 |
| 42 | BIS-TRIS propane | 0.45 | 0.11 | 2.08 | 0.53 |
| 43 | Folic acid | 0.43 | 0.13 | 1.98 | 0.60 |
| 44 | Sodium phosphate dibasic | 0.42 | 0.28 | 1.95 | 1.31 |
| 45 | Pentetic Acid | 0.42 | 0.19 | 1.95 | 0.90 |
| 46 | Potassium chloride | 0.41 | 0.22 | 1.89 | 1.04 |
| 47 | Valine | 0.41 | 0.37 | 1.89 | 1.71 |
| 48 | Isoleucine | 0.41 | 0.21 | 1.88 | 0.97 |
| 49 | Sodium Salicylate | 0.40 | 0.24 | 1.84 | 1.10 |
| 50 | Chlorobutanol Hemihydrate | 0.39 | 0.17 | 1.83 | 0.81 |
| 51 | Methionine | 0.39 | 0.34 | 1.81 | 1.58 |
| 52 | Sodium deoxycholate | 0.38 | 0.12 | 1.75 | 0.54 |
| 53 | TES | 0.38 | 0.28 | 1.75 | 1.30 |
| 54 | PF-68 | 0.38 | 0.27 | 1.75 | 1.23 |
| 55 | Hydroxypropyl Betadex | 0.37 | 0.24 | 1.74 | 1.12 |
| 56 | Potassium phosphate tribasic | 0.37 | 0.18 | 1.71 | 0.85 |
| 57 | Benzethonium Chloride | 0.36 | 0.28 | 1.68 | 1.31 |
| 58 | BIS-TRIS | 0.36 | 0.30 | 1.67 | 1.39 |
| 59 | L-Phenylalanine | 0.36 | 0.23 | 1.66 | 1.06 |
| 60 | y-Aminobutyric Acid | 0.35 | 0.24 | 1.63 | 1.11 |
| 61 | Potassium Gluconate | 0.35 | 0.19 | 1.61 | 0.86 |
| 62 | Sodium Benzoate | 0.34 | 0.25 | 1.59 | 1.17 |
| 63 | Betadex Sulfobutyl Ether Sodium | 0.34 | 0.14 | 1.56 | 0.65 |
| 64 | D-(+)-Trehalose | 0.33 | 0.18 | 1.55 | 0.85 |
| 65 | Asparagine monohydrate | 0.33 | 0.20 | 1.55 | 0.92 |
| 66 | L-Tryptophan | 0.33 | 0.16 | 1.54 | 0.72 |
| 67 | L-(+)-Arabinose | 0.33 | 0.20 | 1.52 | 0.91 |
| 68 | Proline | 0.32 | 0.14 | 1.48 | 0.64 |
| 69 | Polyethylene glycol (3400 Da) | 0.32 | 0.14 | 1.47 | 0.66 |
| 70 | D-Penicillamine | 0.31 | 0.20 | 1.42 | 0.95 |
| 71 | L-Histidine | 0.30 | 0.14 | 1.41 | 0.67 |
| 72 | Potassium phosphate monobasic | 0.30 | 0.12 | 1.41 | 0.57 |
| 73 | B-Cyclodextrin | 0.30 | 0.01 | 1.40 | 0.06 |
| 74 | Polyethylene glycol (1000 Da) | 0.30 | 0.03 | 1.39 | 0.14 |
| 75 | Phenol | 0.30 | 0.25 | 1.38 | 1.17 |
| 76 | Lactose | 0.30 | 0.14 | 1.38 | 0.64 |
| 77 | MOPSO | 0.29 | 0.21 | 1.37 | 0.95 |
| 78 | Lactose Monohydrate | 0.29 | 0.25 | 1.37 | 1.16 |
| 79 | Lactobionic Acid | 0.29 | 0.13 | 1.35 | 0.61 |
| 80 | Beta-Alanine | 0.29 | 0.17 | 1.34 | 0.81 |
| 81 | Methylcellulose | 0.29 | 0.09 | 1.33 | 0.43 |
| 82 | MES | 0.29 | 0.18 | 1.32 | 0.85 |
| 83 | Sucrose | 0.28 | 0.23 | 1.32 | 1.08 |

| | | | | | |
|---|---|---|---|---|---|
| 84 | L-leucine | 0.28 | 0.22 | 1.32 | 1.00 |
| 85 | Sodium Phosphate Monobasic | 0.28 | 0.28 | 1.31 | 1.32 |
| 86 | Ammonium chloride | 0.28 | 0.13 | 1.31 | 0.61 |
| 87 | y-Cyclodextrin | 0.28 | 0.05 | 1.30 | 0.23 |
| 88 | DIPSO | 0.28 | 0.18 | 1.28 | 0.84 |
| 89 | Methyl-B-Cyclodextrin | 0.27 | 0.16 | 1.27 | 0.75 |
| 90 | Saccharin Sodium | 0.26 | 0.19 | 1.21 | 0.86 |
| 91 | 3-(Triphenylphosphonio)propane-1-sulfonate | 0.26 | 0.15 | 1.21 | 0.68 |
| 92 | Niacinamide | 0.25 | 0.16 | 1.17 | 0.75 |
| 93 | L-ascorbic acid | 0.25 | 0.14 | 1.17 | 0.65 |
| 94 | Dextran | 0.25 | 0.09 | 1.16 | 0.42 |
| 95 | D-Sorbitol | 0.25 | 0.12 | 1.16 | 0.55 |
| 96 | Octyl sulfobetaine | 0.25 | 0.14 | 1.15 | 0.65 |
| 97 | HEPES | 0.25 | 0.17 | 1.15 | 0.78 |
| 98 | PF-127 | 0.25 | 0.02 | 1.14 | 0.11 |
| 99 | ACES | 0.24 | 0.15 | 1.12 | 0.68 |
| 100 | Maltose | 0.24 | 0.03 | 1.11 | 0.13 |
| 101 | Betaine | 0.24 | 0.03 | 1.11 | 0.16 |
| 102 | Taurine | 0.24 | 0.15 | 1.11 | 0.69 |
| 103 | Sodium Ascorbate | 0.24 | 0.13 | 1.10 | 0.58 |
| 104 | Dextrose Monohydrate | 0.24 | 0.15 | 1.09 | 0.71 |
| 105 | D-(−)-Fructose | 0.23 | 0.11 | 1.09 | 0.50 |
| 106 | D-glucose | 0.23 | 0.22 | 1.08 | 1.01 |
| 107 | Potassium Acetate | 0.23 | 0.05 | 1.07 | 0.23 |
| 108 | Glycine | 0.22 | 0.12 | 1.04 | 0.57 |
| 109 | Trisodium citrate dihydrate | 0.22 | 0.08 | 1.02 | 0.37 |
| 110 | 3-Amino-4-methoxybenzenesulfonic acid | 0.22 | 0.07 | 1.00 | 0.33 |
| 111 | Phosphate buffered saline (pH 7.4) | 0.22 | 0.09 | 1.00 | 0.41 |
| 112 | Boric Acid | 0.21 | 0.09 | 1.00 | 0.44 |
| 113 | Ammonium Acetate | 0.20 | 0.08 | 0.95 | 0.35 |
| 114 | L-Cysteine hydrochloride | 0.20 | 0.10 | 0.93 | 0.47 |
| 115 | Creatinine | 0.19 | 0.01 | 0.89 | 0.06 |
| 116 | Polyvinylpyrrolidone | 0.19 | 0.06 | 0.88 | 0.27 |
| 117 | Hydrolyzed Soy Protein (2000 Da) | 0.19 | 0.20 | 0.86 | 0.94 |
| 118 | Methylparaben | 0.17 | 0.03 | 0.81 | 0.12 |
| 119 | Hydroxypropyl Cellulose | 0.17 | 0.13 | 0.80 | 0.60 |
| 120 | MOPS | 0.17 | 0.03 | 0.80 | 0.13 |
| 121 | EPPS | 0.17 | 0.16 | 0.79 | 0.74 |
| 122 | Sodium Gluconate | 0.15 | 0.12 | 0.69 | 0.57 |
| 123 | Gentisic Acid | 0.13 | 0.02 | 0.62 | 0.08 |
| 124 | L-Cysteine | 0.13 | 0.04 | 0.60 | 0.20 |
| 125 | Succinic Acid | 0.13 | 0.08 | 0.59 | 0.38 |
| 126 | Gluconolactone | 0.12 | 0.01 | 0.58 | 0.06 |
| 127 | Alpha cyclodextrin | 0.12 | 0.11 | 0.55 | 0.51 |
| 128 | Sodium Bisulfite | 0.11 | 0.02 | 0.53 | 0.08 |
| 129 | Carboxymethylcellulose Sodium | 0.11 | 0.00 | 0.53 | 0.01 |
| 130 | Lauryl sulfobetaine | 0.11 | 0.09 | 0.52 | 0.41 |
| 131 | Tartaric Acid | 0.08 | 0.01 | 0.37 | 0.05 |
| 132 | Malic Acid | 0.08 | 0.01 | 0.35 | 0.06 |
| 133 | Maleic Acid | 0.07 | 0.02 | 0.34 | 0.07 |
| 134 | Glycolic acid | 0.06 | 0.02 | 0.28 | 0.12 |
| 135 | ADA | 0.06 | 0.00 | 0.26 | 0.01 |
| 136 | Citric Acid | 0.04 | 0.01 | 0.20 | 0.03 |
| 137 | Adipic acid | 0.04 | 0.00 | 0.17 | 0.01 |
| 138 | Sodium Sulfite | 0.04 | 0.02 | 0.16 | 0.12 |
| 139 | Sodium Pyrophosphate | 0.03 | 0.01 | 0.13 | 0.04 |
| 140 | Tannic Acid | 0.02 | 0.00 | 0.08 | 0.02 |
| 141 | Carrageenan | 0.00 | 0.00 | 0.01 | 0.01 |
